# A domain-resolution map of *in vivo* DNA binding reveals the regulatory consequences of somatic mutations in zinc finger transcription factors

**DOI:** 10.1101/630756

**Authors:** Berat Dogan, Senthilkumar Kailasam, Aldo Hernández Corchado, Naghmeh Nikpoor, Hamed S. Najafabadi

**Author notes:** These authors contributed equally to this work (Berat Dogan, Senthilkumar Kailasam). Correspondence should be addressed to: H.S.N.

## Abstract

Multi-zinc finger proteins constitute the largest class of human transcription factors. Their DNA-binding specificity is usually encoded by a subset of their tandem Cys2His2 zinc finger (ZF) domains – the subset that binds to DNA, however, is often unknown. Here, by combining a context-aware machine-learning-based model of DNA recognition with *in vivo* binding data, we characterize the sequence preferences and the ZF subset that is responsible for DNA binding in 209 human multi-ZF proteins. We show that *in vivo* DNA binding is primarily driven by ∼50% of the ZFs – these DNA-binding ZFs are under strong selective pressure within and across species, and their mutations affect the expression of hundreds of genes as revealed by pan-cancer trans-eQTL analysis across 18 tissues. Among the genes affected by mutations in multi-ZF proteins, we identify several oncogenic factors regulated by SP1, and show that SP1 up-regulation in cancer promotes the expression of these genes while mutations in SP1 ZFs lead to their repression. Together, these analyses suggest that mutations in DNA-binding ZFs have distinct and widespread regulatory consequences that contribute to transcriptome remodelling in cancer.

## INTRODUCTION

Cys2His2 zinc finger proteins (ZFPs) make up the largest class of human transcription factors (TFs): the human genome encodes ∼750 ZFPs, which constitute ∼45% of all human TFs. Most ZFPs recognize distinct DNA sequences, forming the most diverse regulatory lexicon of all human TFs (Ambrosini et al., 2020; Najafabadi et al., 2015b). These proteins are characterized by the presence of multiple DNA-binding domains known as Cys2His2 zinc fingers (ZFs). Each ZF typically interacts with three to four nucleotides (Wolfe et al., 2000), and the amino acid-base interactions of consecutive ZFs determine the overall DNA sequence specificity of each ZFP.

The human ZFPs on average contain ∼10 ZFs (Najafabadi et al., 2015b), which would correspond to a binding footprint of ∼30 nucleotides. However, the binding sites of ZFPs are often much shorter (Barazandeh et al., 2018), suggesting that only a fraction of the ZFs interact with the DNA while other ZFs might be involved in other functions such as mediating protein-protein (Brayer and Segal, 2008) or protein-RNA (Brown, 2005) interactions. Currently, the DNA-engaging ZFs of only a small subset of ZFPs have been characterized (Baxley et al., 2017; Han et al., 2016; Nakahashi et al., 2013), preventing a comprehensive functional stratification of ZFs. This limited knowledge of the *in vivo* function of each individual ZF hampers our ability to interpret the functional consequences of genetic variations and mutations in ZFs and to understand their role in shaping the cellular and organismal phenotypes.

The modularity of DNA-binding by ZFs provides an opportunity to identify the ZFs that contribute to *in vivo* DNA recognition, based on comparison of the DNA sequence preference of individual ZFs to the sequences of the *in vivo* binding sites of the full length ZFP. However, the sequence preferences of only a fraction of human ZF domains have been experimentally characterized. As an alternative to experimental approaches, several recent studies have developed computational “recognition codes” that predict the binding preference of each individual ZF based on its amino acid sequence (Gupta et al., 2014; Najafabadi et al., 2015b; Persikov et al., 2009; Persikov et al., 2015). These recognition models, however, have limited accuracy for prediction of *in vivo* DNA binding preferences, given that they are primarily derived from *in vitro* data in which individual ZFs are tested in an unnatural context [e.g. as fusion to ZFs of other proteins (Garton et al., 2015; Najafabadi et al., 2015b; Persikov et al., 2015)]. Furthermore, most recognition models are based on the identity of only “four specificity residues”, ignoring the potential contribution of other ZF positions. As a result, the predictions made by existing recognition codes only partially match the observed *in vivo* preferences of ZFPs (Garton et al., 2015; Najafabadi et al., 2015b), limiting their ability to identify the ZFs that contribute to *in vivo* DNA binding.

Here, we take advantage of a large set of recently published *in vivo* ZFP binding preferences (Imbeault et al., 2017; Schmitges et al., 2016) and combine them with *in vitro* data (Najafabadi et al., 2015b) to derive a recognition code of ZF-DNA interaction that is significantly more accurate than existing models. We show that this code captures the contribution of non-canonical ZF residues as well as adjacent ZFs to DNA binding preference, providing an amino acid-resolution map of ZF-DNA interaction. Furthermore, by combining this recognition code with ChIP-seq and ChIP-exo binding data, we identify the human ZF domains that engage with DNA *in vivo*, enabling us to examine the evolutionary pressures acting on these ZFs across species and across individuals in the human population. We show that, in contrast to the ZFs that do not contribute to *in vivo* DNA binding, DNA-binding ZFs are conserved across species and depleted of genetic variation in human. Furthermore, by analyzing the transcriptome data across 18 cancer types, we identify the genome-wide expression changes associated with somatic mutations in ZFs, enabling us to probe the regulatory consequences of disruption of the DNA binding ability of each ZFP and characterize their cancer-associated regulatory programs.

## RESULTS

### A context-aware model of DNA recognition by C2H2-ZFs

The canonical model of C2H2-ZF interaction with DNA primarily includes four “specificity residues”, each of which interacts with four specific sites on the DNA (**Fig. 1a**). However, this model is obtained from a limited number of protein-DNA complex structures, mutation experiments, and *in vitro* data, with other studies suggesting the potential role of other residues in DNA sequence recognition (Persikov and Singh, 2011). In order to examine whether other ZF residues might contribute to sequence-specificity, we correlated the protein sequence of 836 ZF domains from 157 human C2H2-ZF proteins with their *in vivo* binding preferences, which revealed highly significant dependencies between amino acid identity and DNA base identity for all expected canonical interactions (**Fig. 1b**). The significant associations, however, were not limited to these positions, and included other interactions, e.g. between residue +2 of the ZF and DNA triplet position +3, and residue +6 of the ZF and all three DNA triplet positions as well as DNA position +3 of the upstream triplet (**Fig. 1b**). Interestingly, many of the associations observed from analysis of *in vivo* binding preferences can also be captured as hydrogen bonds and other non-covalent interactions by molecular dynamics (MD) analysis of the structure of the Egr1 zinc finger protein in complex with its cognate DNA (**Fig. 1c**). Example structures highlighting some of these contacts are shown in **Fig. 1d-e**, including a non-canonical hydrogen bond and a π-π stacking interaction between DNA and positions +2 and +6 of ZF1, respectively.

**Figure 1.**
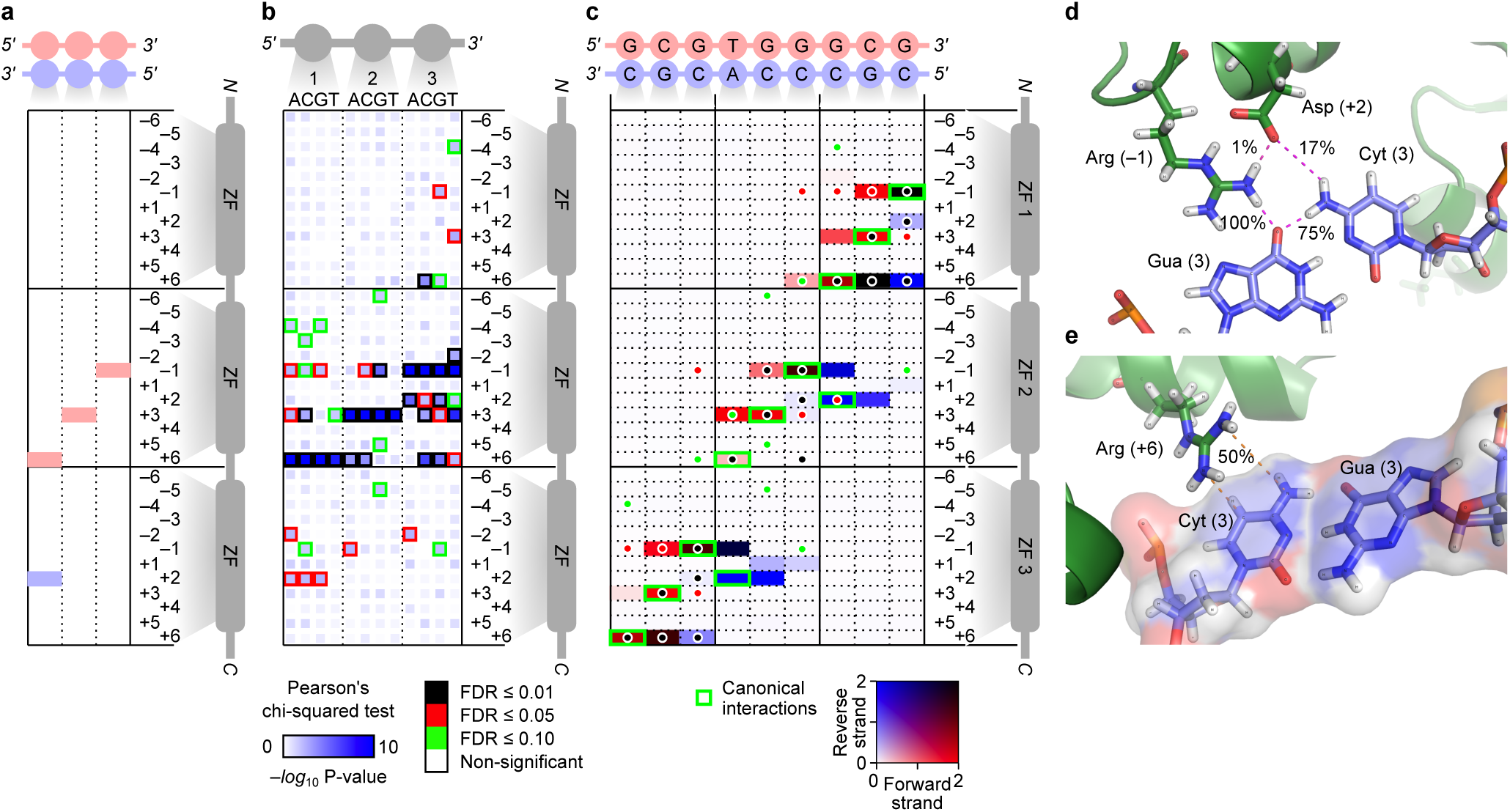
The canonical model of C2H2-DNA interaction only explains a subset of possible contacts. (**a**) Schematic representation of the canonical interactions between ZF positions –1, +2, +3, and +6 with a DNA triplet. Red and blue depict interaction with forward and reverse strand, respectively. Residue numbers are relative to the start of ZF alpha helix. (**b**) Associations between the identity of different ZF residues and each of the four bases at different DNA positions, based on *in vivo* motifs (Imbeault et al., 2017; Schmitges et al., 2016). We divided the *de novo*-identified *in vivo* motifs into three-nucleotide partitions (triplets) each representing the binding preference of one ZF domain, and examined whether the amino acid identity at each ZF position, including those of the N-terminal and C-terminal neighbors of the directly contacting ZF, is informative about the base preference at each triplet position (chi-square test after dichotomizing the affinity values for each base at each position). The color gradient represents the P-value of the chi-square test; the border color represents FDR. Data underlying this figure are included in **Table S1**. (**c**) A heatmap showing the average number of all-atom contacts between each residue of Egr1 and the nucleotide bases of the forward and reverse strand of the target DNA, based on a 1μs-long MD simulation of PDB structure 4×9J. The color gradient stands for the average number of contacts observed per frame of MD simulations, with red and blue representing contact with forward and reverse DNA strands, respectively. The dots represent significant associations based on association analysis of *in vivo* binding preferences, with the dot colour mirroring the border colour in panel b. Underlying data are provided in **Table S2**. (**d**) Structural illustration of a non-canonical hydrogen bond between position +2 of Egr1 ZNF1 and the reverse strand base at position 9 of the DNA sequence (i.e. position 3 of the third triplet), which can be observed at ∼17% occupancy. (**e**) The planar guanidino group of Arginine in position +6 of ZF1 forms a π-π stacking interaction with reverse strand base of the DNA position 9 (∼50% occupancy).

These analyses suggest that amino acid-base interactions are not limited to the canonical model, and non-canonical interactions, including those mediated by adjacent ZFs, can be captured by analysis of *in vivo* binding preferences of ZFs. The available *in vitro* data (Najafabadi et al., 2015b), on the other hand, provide a higher coverage and larger training dataset for modeling ZF-DNA interactions. Therefore, we sought to obtain a *compound recognition code* (hereafter referred to as C-RC), which combines *in vitro* and *in vivo* data in order to obtain optimal accuracy for prediction of the DNA sequence preference of ZFs given their amino acid sequences. We performed a systematic feature selection to identify the ZF residues that maximize prediction accuracy when used as input to random forest regression models (Breiman, 2001) that predict specificity for each of the four bases at each DNA triplet position (**Fig. 2a**). We identified different features that optimize prediction accuracy in different contexts – the contexts that we considered included whether the ZF was located at the N-terminus, C-terminus, or the middle of a DNA-engaging ZF-array (**Fig. 2a**). We also found that encoding the amino acids based on a low-dimensional representation of their biochemical and structural properties [as opposed to one-hot encoding (Najafabadi et al., 2015b)] substantially reduces the complexity of the model (**Fig. S1a-b**) and increases model accuracy and generalizability (**Fig. S1c**).

**Figure 2.**
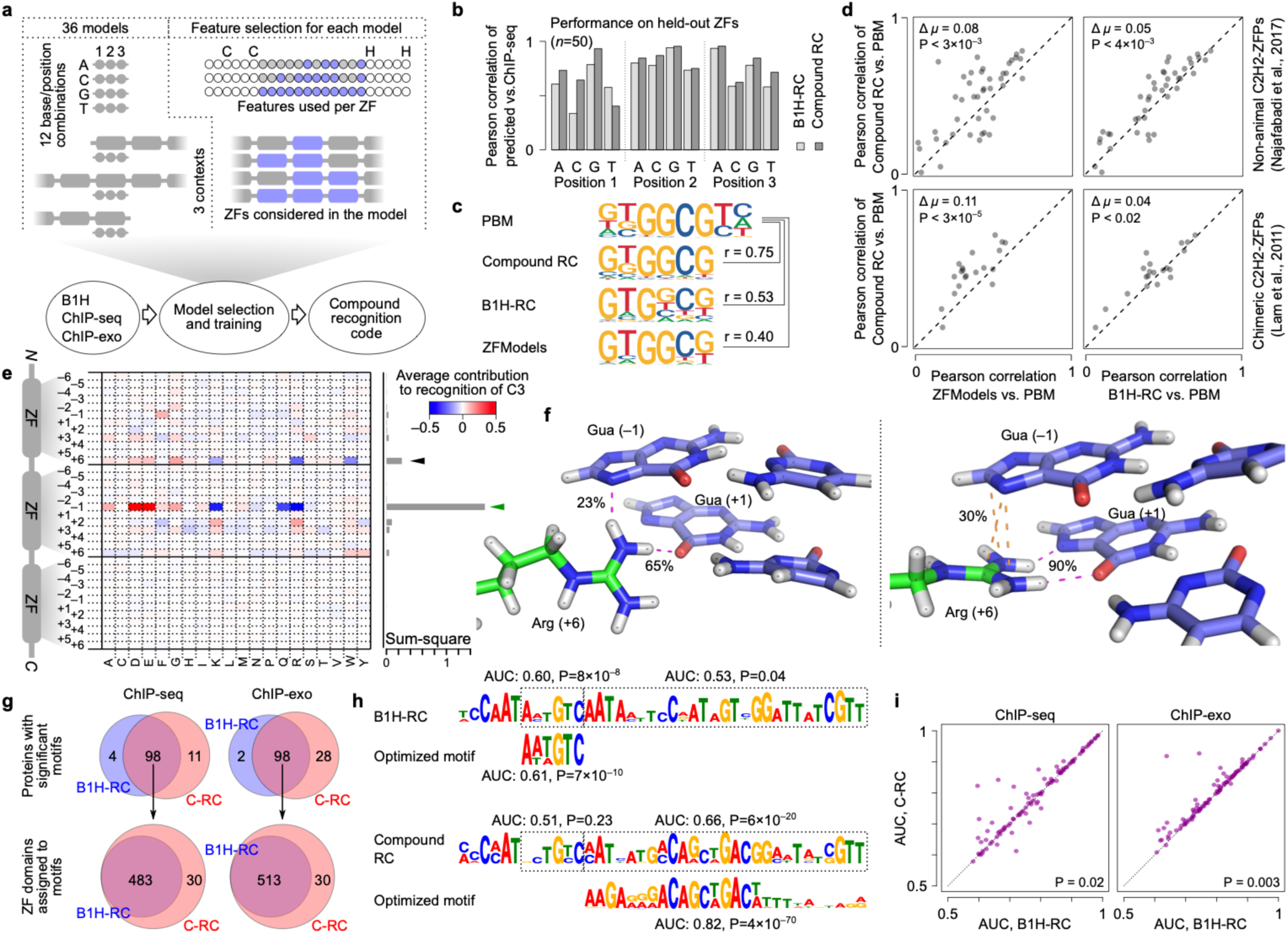
Construction and evaluation of the compound recognition code (C-RC). (**a**) Schematic representation of the procedure for training C-RC. Thirty-six models were trained separately (top left), corresponding to recognition of four bases at three DNA positions in three different contexts. Feature selection for each model included selection of residues to be considered for each ZF, as well as the choice of ZFs (top right). See **Fig. S2** for detailed representation of the selected features for each model. (**b**) Comparison of the performance of C-RC and a B1H-based recognition code (Najafabadi et al., 2015b) (B1H-RC) for predicting the base probabilities at different triplet positions for 50 ChIP-seq ZFs that were excluded from the feature selection procedure and model training. (**c**) Comparison of the motifs predicted by C-RC, B1H recognition code, and ZFModels (Gupta et al., 2014) vs. the motifs obtained by PBM (Najafabadi et al., 2017) for a ZFP from *Rhizopus delemar* (UniProt ID I1BM86), as an example. Pearson correlation similarities are calculated by MoSBAT-e (Lambert et al., 2016). (**d**) Systematic comparison of C-RC, B1H-RC, and ZFModels predictions vs. PBM motifs from non-animal ZFPs (Najafabadi et al., 2017) and chimeric ZFP constructs (Lam et al., 2011). Pearson correlation similarity of motif pairs are calculated by MoSBAT-e (Lambert et al., 2016). P-values are based on two-tailed paired t-test. (**e**) Representation of the rules encoded in the C-RC for recognition of base C at position 1 of the DNA triplet (see **Fig. S3a** for details). The heatmap shows the average contribution of each amino acid at different ZF positions. The bar graph on the right shows the squared sum of values at each ZF position. The canonical specificity residue that is known to interact with this position of DNA is denoted using the green arrow, while the black arrow represents a potential new interaction with position +6 of the N-terminal neighboring ZF. See **Fig. S3a** for recognition of other bases at different triplet positions. (**f**) The Arginine at position +6 of ZF4 forms a non-canonical interaction with position –1 of its cognate DNA triplet (i.e. position +3 of the preceding triplet), based on MD simulations of CTCF in complex with its target DNA. The interaction exists in two dominant conformations: a hydrogen bond (23% of MD simulation frames) between the Arginine and the base (left) and a stacking overlap (30% of MD simulation frames) between Arginine and the guanidino group and the DNA base (right). (**g**) Top: systematic comparison between B1H-RC and C-RC for the number of motifs that are significantly enriched in ChIP-seq data (Schmitges et al., 2016) (left) or ChIP-exo data (Imbeault et al., 2017) (right). The Venn diagrams correspond to cases in which the C-RC prediction was enriched in peaks relative to dinucleotide shuffled sequences (hypergeometric test, P<0.0001), and the final optimized motif was still significantly similar to the seed C-RC motif (P<0.001). Bottom: for motifs that are common between B1H-RC and C-RC, the number of associated ZFs are compared. (**h**) Comparison of the B1H-RC-predicted motif (top) and C-RC-predicted motif (bottom) along with their ChIP-seq-based optimized versions for ZNF317, as an example. (**i**) For proteins with significant motifs, the AUROC values of the optimized versions are compared between B1H-RC and C-RC for the ChIP-seq dataset (left) and ChIP-exo dataset (right). P-values are based on two-tailed t-test.

The final recognition code consists of 36 random forests: 12 random forests for each of the N-terminal, C-terminal, or middle ZF contexts, with each random forest predicting the preference for binding to one of the four possible bases at one of the three DNA triplet positions (**Fig. S2**). We benchmarked the performance of C-RC using different evaluation schemes. First, we used 50 randomly selected ZF-target pairs from ChIP-seq data, which were excluded from all stages of the random forest training, including feature selection. As shown in **Fig. 2b**, C-RC outperformed a previous B1H-based recognition code (B1H-RC) (Najafabadi et al., 2015b) in predicting the base preferences for 11 out of 12 base-position combinations. Next, we compared the accuracy of C-RC with B1H-RC (Najafabadi et al., 2015b) and another recent recognition code, ZFModels (Gupta et al., 2014), using an independent protein-binding microarray (PBM) compendium from a set of non-animal ZFPs (Najafabadi et al., 2017) and a set of synthetic ZFPs created from concatenating different ZFs from different proteins (Lam et al., 2011). C-RC produced motifs that were significantly more similar to the PBM motifs than both B1H-RC and ZFModels predictions in both datasets (**Fig. 2c-e**), suggesting that the new code outperforms the state-of-the-art irrespective of the nature of the reference data (*in vivo* or *in vitro*) or the source of the protein (human, non-animal, or synthetic). We found that C-RC encodes the canonical ZF-DNA interactions (**Fig. S3a)** as well as strong associations that are not part of the canonical model, most prominently between position +3 of the DNA triplet and residue +6 of the N-terminal neighboring ZF (**Fig. 2e**). MD simulations of the CCCTC-binding factor (CTCF) in complex with its cognate DNA indeed identified a non-canonical interaction between position +3 of the DNA triplet that is adjacent to ZF5 and residue +6 of ZF4 (**Fig. S3b** and **Fig. 2f**), supporting the biological relevance of this non-canonical interaction. This interaction was also observed in MD simulations of the Egr1-DNA complex, between position +3 of the second DNA triplet and residue +6 of ZF1 (**Fig. 1c**). To further demonstrate that the C-RC faithfully encodes ZF-DNA interactions, we performed an *in silico* mutation screen using C-RC to predict the effect of various CTCF mutations on its DNA binding affinity. This analysis revealed a quantitative relationship between the C-RC-predicted mutation effects and the strength of H-bonds observed in MD simulations (**Fig. S3c-d**). Together, these analyses suggest that C-RC outperforms existing C2H2-ZF recognition models, encodes canonical as well as non-canonical interactions between ZF residues and DNA, and can predict these interactions quantitively.

### A domain-resolution map of DNA binding by ZFPs reveals function-dependent conservation of ZFs

Human ZFPs on average contain ∼10 ZFs per protein (Najafabadi et al., 2015b), which is substantially longer than what would be expected from the binding preference of most transcription factors, suggesting that only a fraction of the ZFs of most proteins engage with DNA. Combining *in vivo* binding data with the predicted DNA preferences of the ZFs enables the identification of the ZF domains that engage with DNA (Najafabadi et al., 2015a). We therefore sought to combine our new recognition code with previously published ChIP-seq and ChIP-exo data of ZFPs (Imbeault et al., 2017; Schmitges et al., 2016) to characterize their *in vivo* binding preferences and the ZF domains that engage with the DNA. We used our previously described motif discovery framework (Najafabadi et al., 2015a), which starts from the predicted binding preferences of all possible ZF arrays and identifies the ZF array whose binding preference maximally explains the observed *in vivo* binding sequences, along with its optimized binding motif. We observed a substantial improvement in the ability of this framework to identify *in vivo*-enriched motifs when it starts from the C-RC predictions instead of B1H-RC predictions (7% increased success rate for ChIP-seq and 26% increased success rate for ChIP-exo; **Fig. 2g**). Furthermore, for cases where both B1H-RC and C-RC produced *in vivo*-enriched motifs, C-RC was able to identify a larger number of ZFs that engage with DNA (**Fig. 2g-h**), with optimized motifs that had higher quality (**Fig. 2i**).

Overall, we identified significant motifs for 109 ChIP-seq and 126 ChIP-exo datasets (**Fig. 3a** and **S4**, respectively), encompassing a total of 209 unique proteins. On average, about 56% of the ZFs of each protein appear to engage with DNA *in vivo* in the ChIP-seq dataset and 45% in the ChIP-exo dataset, although these ZFs vary substantially in terms of their sequence specificity, as measured by the information content of the 3-nucleotide motif that corresponds to each ZF (**Fig. 3a** and **S4**). Among the 209 proteins, we identified four ZFPs whose DNA-binding domains were previously characterized (CTCF, PRDM9, OSR2, and GTF3A), all of which were in agreement with our functional ZF stratification (**Fig. S5a**). This observation, although limited, supports the notion that by combining C-RC predictions and *in vivo* data we can identify the ZFs that engage with DNA. A more comprehensive source of evidence comes from the observation that CRC-predicted DNA-engaging ZFs are overall more conserved across vertebrates than non-binding ZFs or ZFs that bind to DNA with low sequence specificity (**Fig. 3b-c**). This trend can be seen when we aggregate all ZFs from all ZFPs that have ChIP-seq data (**Fig. 3b**), as well as when we analyze the ZF domains of each ZFP individually (**Fig. 3c**; we focused on the ChIP-seq dataset since the ChIP-exo dataset is only limited to KRAB-containing proteins). We also separately analyzed the conservation pattern of ZF domains in ZFPs that contain a KRAB domain – these ZFPs are relatively recent and are often involved in repression of transposable elements (Imbeault et al., 2017; Jacobs et al., 2014; Najafabadi et al., 2015b; Schmitges et al., 2016), and their role in regulating host gene expression is largely unknown. Surprisingly, despite their overall lower conservation across vertebrates, likely due to their more recent origin, the correlation between conservation and DNA-binding can be clearly seen in KRAB-containing ZFPs (**Fig. S5b**), suggesting that DNA-binding ZFs are more conserved than non-binding ZFs both in KRAB-containing and non-KRAB ZFPs.

**Figure 3.**
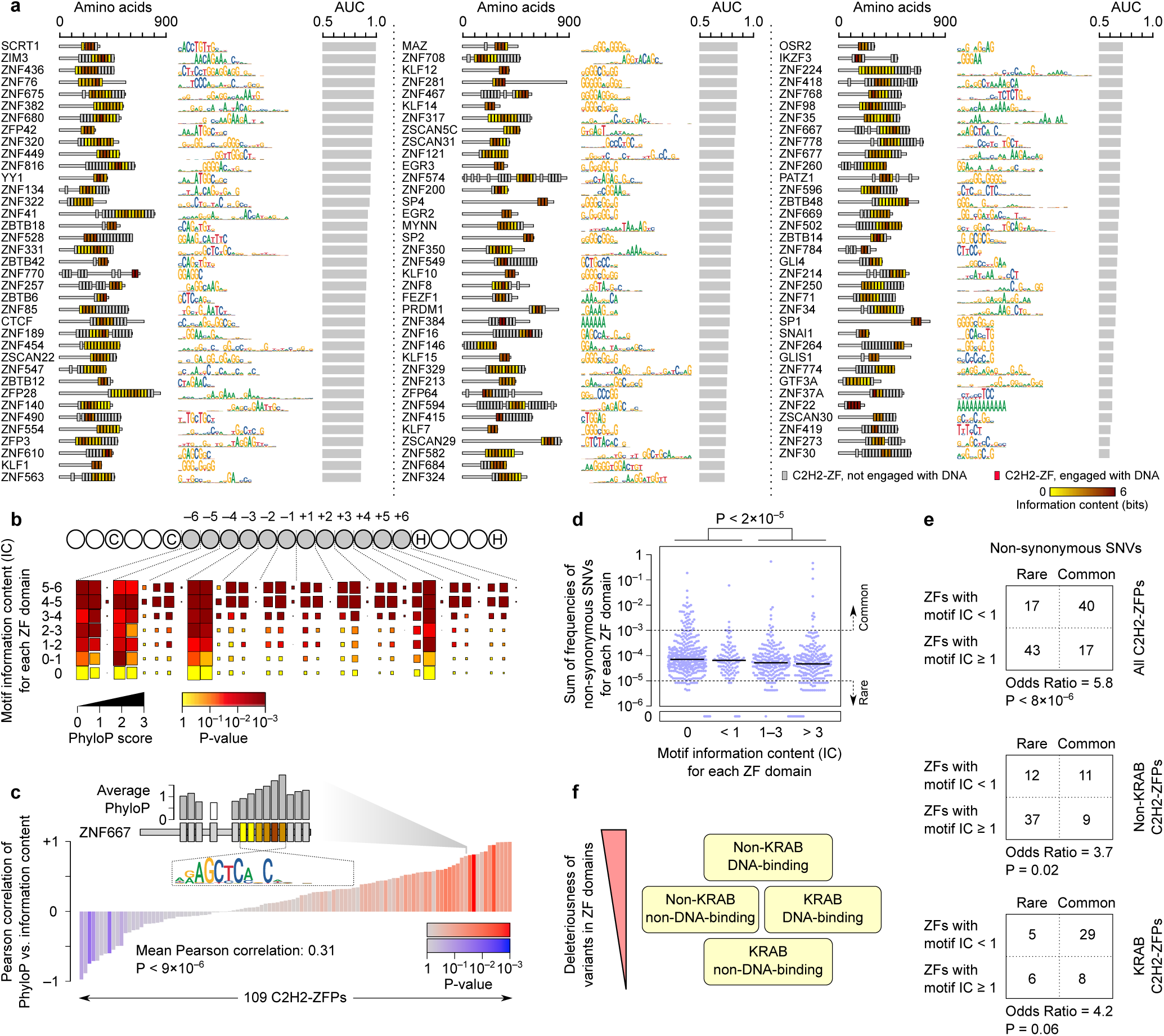
DNA-binding ZFs are under stronger purifying selection for both KRAB and non-KRAB ZFPs. (**a**) Motifs identified by recognition code-assisted analysis of ChIP-seq data for ZFPs. For each protein, the domain structure is shown on the left (only the ZF domains), and the identified motif is shown in the middle. The bar graph shows the AUROC for distinguishing the binding site sequences from dinucleotide-shuffled sequences. The ZFs that correspond to the identified motifs are highlighted, with the colour gradient representing the sequence-specificity of the ZF (as measured by information content). See **Fig. S4** for motifs obtained from ChIP-exo data. Position-specific frequency matrices are provided in **Data Files S1 and S2** for ChIP-seq and ChIP-exo data, respectively. (**b**) PhyloP (Pollard et al., 2010) conservation score profile across ZFs that do not engage with DNA (i.e. IC=0) and those that engage with DNA with varying sequence-specificity. Each set of three columns represents one codon in the ZF-encoding sequence, and each row represents one set of ZFs, binned by the information content (IC) of their associated triplet in the *in vivo* motif. The box size represents the average phyloP score, whereas the color gradient represents the difference of conservation compared to ZFs that do not engage with DNA (P-value based on one-tailed t-test). Data provided in **Table S5a**. (**c**) Correlation of the phyloP score profile of each ZFP with the IC of the ZFs. The inset illustrates an example in which the phyloP score of the ZFs (bar plot above the ZNF667 domain structure) correlates with the IC of the ZF-associated triplets (color of the ZF domains, similar to panel a). For each ZFP, the P-value associated with the Pearson correlation is shown using the color of the associated bar (two-tailed correlation t-test). The overall P-value for deviation of the mean of correlations from zero is also shown (two-tailed t-test). Also see **Fig. S5b-c**. (**d**) Minor allele frequencies of non-synonymous variants for ZFs that do not engage with DNA (IC=0), ZFs with low sequence-specificity (0<IC<1), and ZFs with high sequence-specificity (IC≥1). Each dot represents one ZF, and the y-axis shows the sum of frequencies of all non-synonymous minor alleles that overlap with the ZF. P-value is based on two-tailed t-test. Data provided in **Table S5b**. (**e**) Overlap of rare and common non-synonymous variants with ZFs that engage with DNA with high specificity (IC≥1) and ZFs that do not engage or have low specificity (IC<1). The top table represents all C2H2-ZF proteins with ChIP-seq-based motifs, whereas the next two tables are stratified based on absence or presence of a KRAB domain in the protein. (**f**) A schematic model showing the inferred deleteriousness of variants in different ZFs, as suggested by the frequencies of common and rare variants.

The higher conservation of DNA-engaging ZFs suggests that mutations in these regions are overall more deleterious than mutations in ZFs that do not bind DNA (which may potentially have other functions). We set out to confirm this hypothesis by analysis of population-level genetic variations in human ZFs. Using genetic variation data from gnomAD (Lek et al., 2016), which encompasses variants identified from >140,000 individuals, we observed that missense variations in ZFs that engage with DNA with high specificity are significantly less frequent than missense variations in ZFs that do not engage with DNA *in vivo* or have low sequence specificity (**Fig. 3d**). We also specifically examined the frequency of rare missense variants (minor allele frequency < 10^−5^) and more common variants (minor allele frequency > 0.001), reasoning that extremely rare variants are likely to be recent and, therefore, have not been filtered by negative selection yet, as opposed to common variants (Jaganathan et al., 2019). We noticed that common variants are significantly depleted from DNA-engaging ZFs compared to non-binding or low-specificity ZFs (**Fig. 3e**), suggesting that these ZFs are under stronger negative selection. This trend holds for both KRAB-containing ZFPs and non-KRAB ZFPs, confirming the pattern that we observed by analysis of ZF conservation across vertebrates. Nonetheless, we observed overall stronger depletion of common variants in non-KRAB ZFPs. This trend suggests a stratified model of selective pressure on the ZFs based on their function, in which mutations at the DNA-binding ZFs of non-KRAB proteins are overall the most deleterious ones, whereas mutations at the non-DNA-binding ZFs of KRAB proteins are least affected by negative selection (**Fig. 3f**).

### Somatic mutations in DNA-binding ZFs accompany widespread gene expression remodeling in cancer

Previous reports have suggested that somatic mutations in several cancer types are enriched in critical positions of C2H2 zinc fingers (Munro et al., 2018). Our results suggest that a considerable fraction of the somatic mutations that overlap DNA-binding ZFs should be detrimental to ZF-DNA interaction, given the tight association between conservation and DNA-binding ability across all ZF residues (**Fig. 3b**) and depletion of common genetic variants in DNA-binding ZFs (**Fig. 3d-e**). Therefore, these mutations should have direct and/or indirect regulatory consequences. We studied transcriptome changes associated with ZF somatic mutations across 2538 tumour samples and 18 cancer types (Hoadley et al., 2018) (**Table S6**), in order to understand the gene regulatory consequences of these mutations. For each ZFP, we selected the samples with no copy number alterations (CNAs) at the ZFP locus, and then modeled the association between expression of each query gene and the presence of at least one missense mutation in the DNA-binding zinc fingers of the ZFP, while correcting for the confounding effect of tissue of origin (cancer type), the cancer type-specific effects of sex and age, and the CNAs at the query gene (**Fig. 4a**). This pan-cancer meta-analysis revealed substantial association between genome-wide gene expression and somatic mutations at the DNA-binding ZFs of specific ZFPs (**Fig. 4b, Data File S3**).

**Figure 4.**
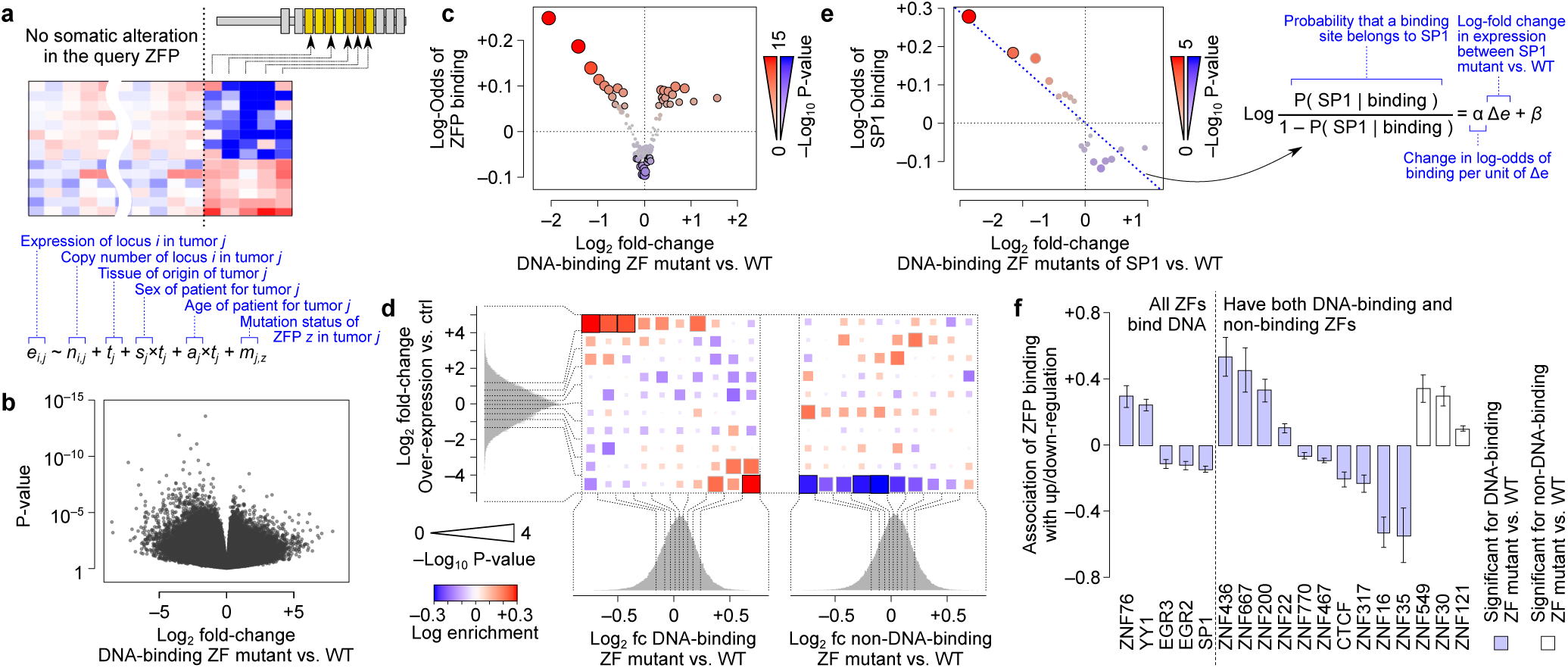
Regulatory consequences of somatic mutations in DNA-binding ZFs. (**a**) Schematic representation of the model used for identifying mutation-associated gene expression changes in cancer. (**b**) Volcano plot for association of gene expression and somatic mutations in DNA-binding ZFs for 1,304,830 gene-ZFP pairs. The x-axis represents the effect size of the somatic mutation, and the y-axis represents the associated P-value. (**c**) Enrichment of ChIP-seq-based ZFP-gene regulatory connections among the ZFP-gene pairs with varying degrees of mutation-expression association. ZFP-gene pairs were grouped into 100 equally populated bins based on the mutation effect size (x-axis) and, in each bin, the enrichment of ZFP-gene pairs that represent a ChIP-seq-based regulatory link relative to expectation was calculated (see **Methods**). The y-axis represents log-odds of regulatory links, each circle represents one bin, and the circle size/color represents P-value (logistic regression). (**d**) For 63 ZFPs that have RNA-seq data from over-expression in HEK293 cells (Schmitges et al., 2016), we binned the ZFP-gene pairs based on the joint distribution of the effect size of ZFP over-expression (y-axis) and the effect size somatic mutations (x-axis) in DNA-binding ZFs (left) or non-DNA-binding ZFs (right). For each bin, the log-odds for ChIP-seq-based ZFP-gene regulatory links relative to expectation is shown using the color of boxes in the heatmap, while the size of the boxes correspond to P-value (logistic regression, see **Methods**). The histograms and the dotted lines represent the distribution of effect sizes and the boundaries of the bins. (**e**) Similar to panel (c) but limited to SP1-gene pairs. Genes are binned into 20 equally populated bins based on the effect size of SP1 mutation. (**f**) ZFPs whose ChIP-seq-based regulatory targets are enriched among genes that are positively or negatively associated with the ZFP somatic mutation. A logistic regression was used to assess the significance of the association between the presence of a peak near gene TSS and the effect size of mutation on gene expression (shown in panel e; also see **Methods**), separately for somatic mutations in DNA-binding ZFs (blue bars) and non-DNA-binding ZFs (white bars). The y-axis shows the regression coefficient, and the error bars represent standard error of mean. Only significant ZFPs (FDR<0.005) are shown.

Several lines of evidence suggest that these ZFP-gene associations reflect bona fide regulatory interactions: First, we observed a significant overlap between the ZFP-gene associations we identified by analysis of somatic mutations and the direct ZFP-gene associations identified by ChIP-seq (**Fig. 4c**). Secondly, this overlap is consistent with disruption of ZFP function, as revealed by analysis of RNA-seq data from over-expression of 63 ZFPs (Schmitges et al., 2016) (**Table S7**): we found that direct ZFP targets (based on ChIP-seq) are enriched among genes that respond in opposite directions to somatic mutations and ZFP over-expression (**Fig. 4d**). In contrast to somatic missense mutations in DNA-binding ZFs, somatic mutations in ZFs that do not bind DNA *in vivo* did not show such a pattern (**Fig. 4d**), suggesting that mutations in DNA-binding ZFs are more likely to have a transcriptomic signature that opposes phenotypic activation of the ZFP. Thirdly, we found that the direct regulatory targets of each ZFP (**Data File S4**) overall show a consistent response to somatic mutations in DNA-binding ZFs (i.e. respond in the same direction). For example, SP1 binding sites are specifically enriched near the transcription start sites (TSS’s) of genes that are down-regulated as a result of SP1 mutation (**Fig. 4e**). Overall, we found 15 ZFPs in which somatic mutation of DNA-binding ZFs leads to a consistent response in their direct targets (FDR < 0.005). In contrast, somatic mutations of non-DNA-binding ZFs showed a similarly significant pattern in only three ZFPs (**Fig. 4f**). Finally, we observed that loss of heterozygosity (LOH) in each ZFP locus leaves a gene expression signature that is highly consistent with that of somatic missense mutations in DNA-binding ZFs (**Fig. 5a**), suggesting that somatic mutations of DNA-binding ZFs, which are often monoallelic, have regulatory effects similar to LOH.

**Figure 5.**
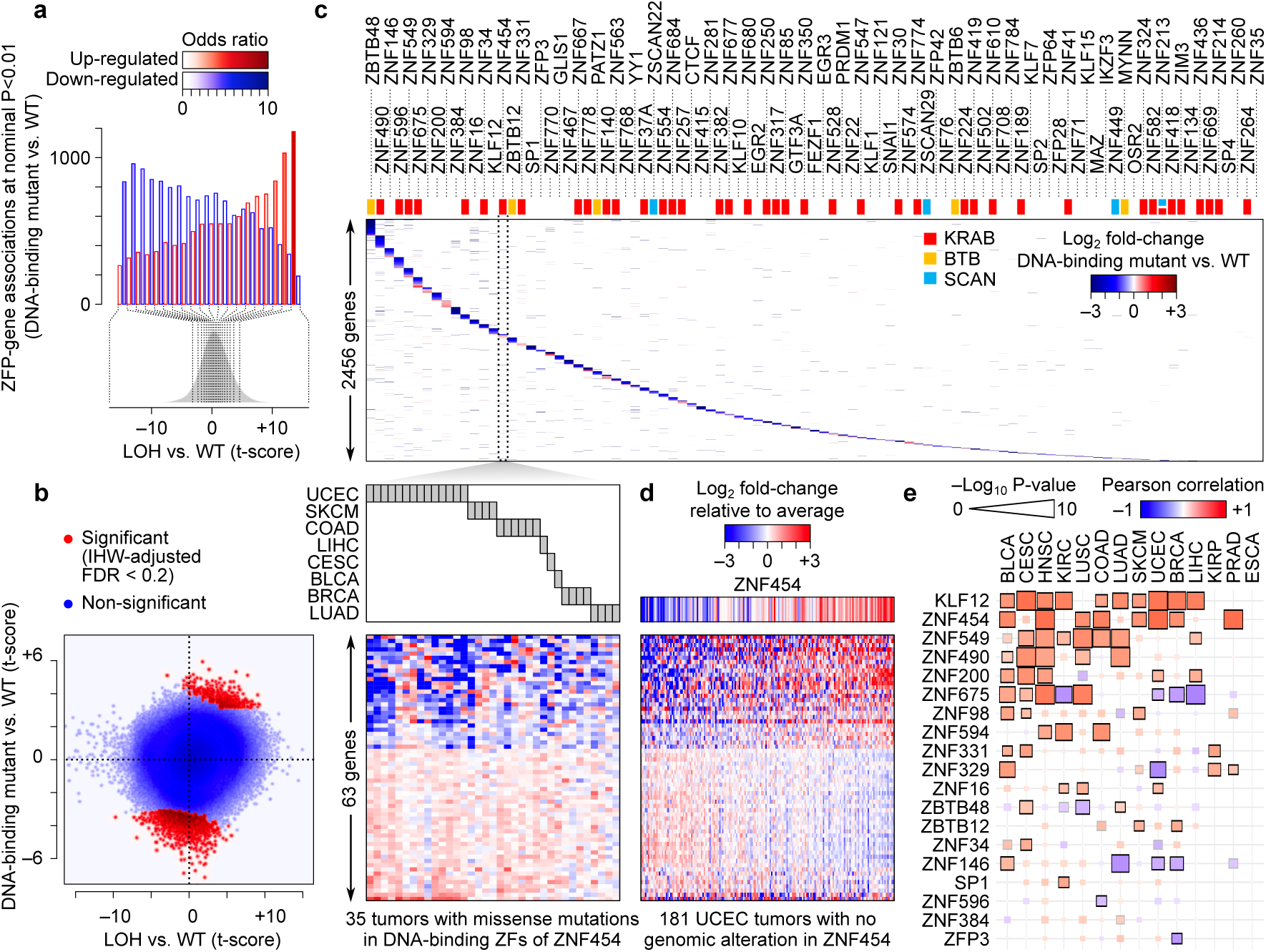
Using somatic alterations to identify gene regulatory targets of ZFPs. (**a**) Consistence between the effect of ZFP somatic mutations and ZFP loss-of-heterogeneity (LOH). ZFP-gene pairs were grouped into 20 bins based on the expression effect size of ZFP LOH (x-axis), and within each bin, the number of ZFP-gene pairs that pass P-value threshold of 0.01 for positive association or negative association with somatic mutation was calculated (y-axis). The color gradient shows the enrichment of ZFP-gene associations with positive mutation effect size (red) or negative mutation effect size (blue). (**b**) Identification of significant associations between ZFP somatic mutation and gene expression, using the association between ZFP LOH and gene expression as a prior for increased statistical power. Each dot represents one ZFP-gene pair, with significant gene pairs (IHW-adjusted FDR<0.2) shown in red. (**c**) Heatmap of significant ZFP-gene associations (top). Each column is one ZFP, each row is one gene, and the colour gradient shows the size of the effect of somatic mutation in the DNA-binding ZFs of each ZFP on the expression of each gene. As an example, genes that are significantly associated with DNA-binding ZF mutations in ZNF454 are shown at the bottom heatmap, with each column representing one mutation-containing tumour sample. The cancer type of each sample is shown on top of the heatmap. (**d**) The correlation between ZNF454 expression across UCEC tumours and expression of its regulon. Each row is a gene, in the same order as the bottom heatmap of panel (c). Each column is one sample, limited to those that do not have any somatic alteration in the ZNF454 locus. The ZNF454 expression itself is shown on top. (**e**) Concordance between ZFP mutation effect size and ZFP-gene co-expression. For each ZFP and each cancer type, the Pearson correlation between the genome-wide effect size of somatic DNA-binding ZF mutation and the ZFP-gene expression correlations were calculated (see **Methods**). The color gradient shows the Pearson correlation, with red depicting correlations that are in the expected direction and blue representing correlations that are opposite of the expected direction. The box size shows the associated P-value (t-test). Significant Pearson correlations are shown with a black border (FDR<0.01) or grey border (FDR<0.05).

### ZFPs regulate hundreds of genes across cancers

The above results enabled us to identify the set of genes whose expression is significantly affected by the missense mutations that disrupt DNA-binding capability of each ZFP, including direct and/or indirect targets of the ZFP. To increase statistical power, we leveraged the LOH-associated gene expression changes (**Fig. 5a**), and used them as a prior to identify mutation-associated expression changes using independent hypothesis weighting (Ignatiadis et al., 2016) (**Fig. 5b**). Overall, we identified 3366 significant ZFP-gene associations (FDR < 0.2) between 2456 genes and 95 ZFPs (**Fig. 5c, Table S8**). The size of the identified ZFP regulons ranged from only 1 gene for ZNF35 to 201 genes for ZBTB48, suggesting varying degrees of power for detecting gene regulatory effects of different ZFPs based on analysis of somatic mutations.

To evaluate their fidelity, we examined the agreement between our mutation-based ZFP-gene associations and the co-expression of ZFP-gene pairs – genes that are dysregulated as a result of a ZFP mutation are expected to also respond to the level of expression of that ZFP, even across samples that have neither CNA nor any mutation at the ZFP locus (an example is shown in **Fig. 5d**). Overall, we observed a trend consistent with this expectation across multiple cancer types: we identified 83 ZFP-cancer pairs for which there was a significant association between mutation-based regulatory links and co-expression. Among these significant associations, ∼81% (67 out of 83, binomial P<10^−8^) had the expected direction, i.e. genes that were down-regulated as a result of ZFP mutation were positively correlated with ZFP expression, and mutation-up-regulated genes were negatively correlated with ZFP expression (**Fig. 5e**). Together, these results support the notion that somatic missense mutations in DNA-binding ZFs disrupt the regulatory function of ZFPs, revealing hundreds of genes that are either directly or indirectly affected by the ZFP activity.

### Mutations in SP1 zinc fingers inhibit its oncogenic regulatory program

Among the ZFPs that showed co-expression with their mutation-responsive genes, the transcription factor SP1 is commonly up-regulated across different cancer types (Beishline and Azizkhan-Clifford, 2015). SP1 over-expression is associated with tumour progression (Hsu et al., 2012) and poor disease outcome (Wang et al., 2003), and its inhibition reduces tumor formation ability of cell lines (Lou et al., 2005). Our somatic mutation analysis revealed that mutations in the DNA-binding domains of SP1 are associated with significant down-regulation of 54 genes (**Table S8**). To further characterize the role of SP1 in regulating the expression of these genes, we analyzed data from cell lines in which SP1 expression was modulated, including CRISPRi-mediated inhibition of SP1 in K562 cells (Consortium, 2012) and over-expression of SP1 in HEK293 cells (Schmitges et al., 2016). We observed that SP1 inhibition leads to significant down-regulation of SP1 mutation-responsive genes [GSEA (Subramanian et al., 2005) P < 0.0006, **Fig. 6a**], while phenotypic activation of SP1 by over-expression leads to their up-regelation (P < 0.0007, **Fig. 6b**). We identified a core set of genes that are down-regulated by SP1 inhibition, up-regulated by SP1 phenotypic activation, and down-regulated by SP1 mutations (**Fig. 6c**) – these genes include CD70, a critical factor in tumour cell survival and immune modulation that is previously reported to be under SP1 control (Riether et al., 2015), and APOBEC3F, a member of the APOBEC family that is implicated in carcinogenic mutagenesis (Roberts et al., 2013). In further support of the role of SP1 in regulation of these genes in cancer, we observed that the majority of them are up-regulated in the cancer types in which SP1 is up-regulated (**Fig. 6d**).

**Figure 6.**
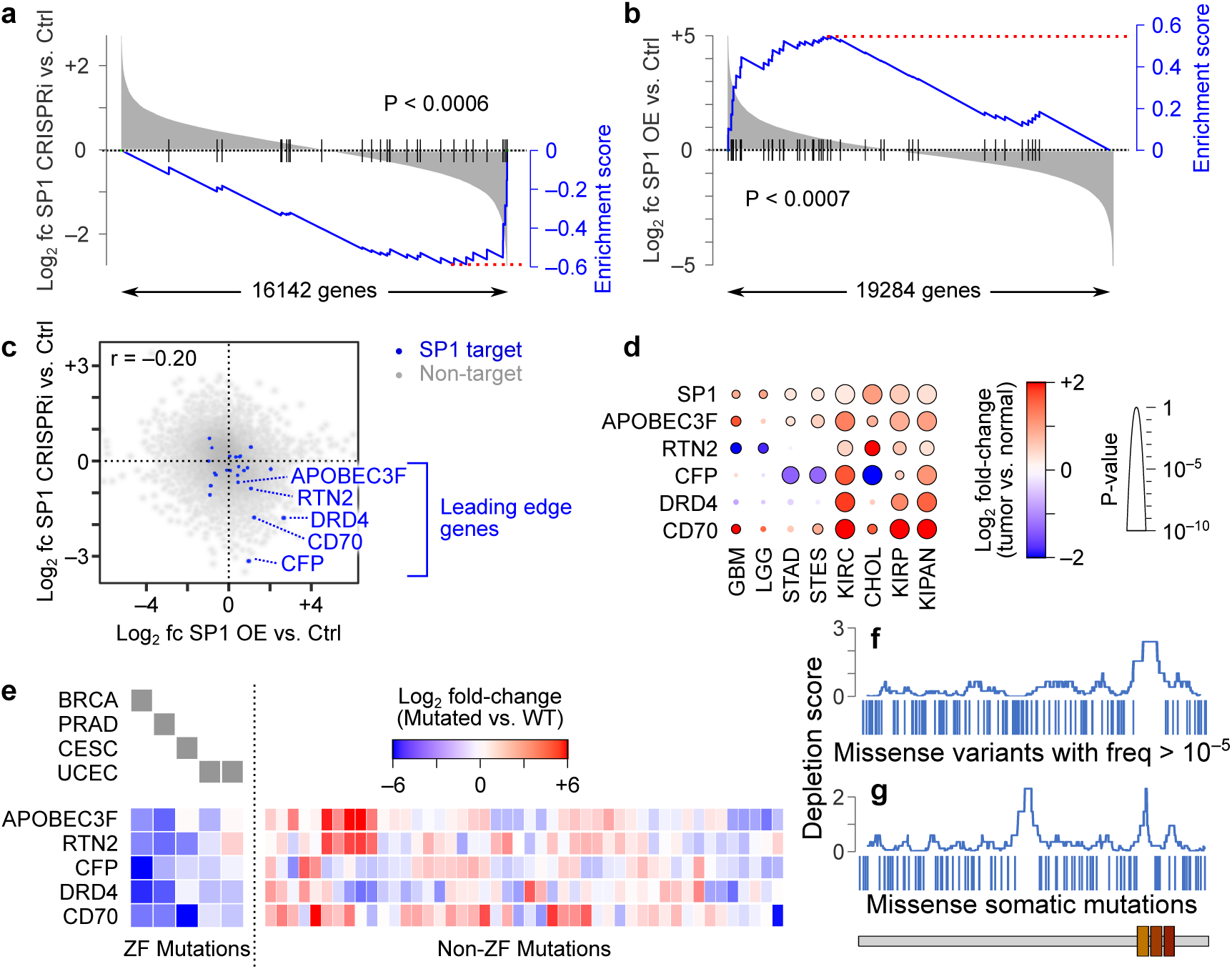
Mutations in SP1 zinc fingers inhibit the SP1-mediated activation of gene expression. (**a**) Gene set enrichment analysis (Subramanian et al., 2005) (GSEA) for SP1 inhibition in K562 cells [data from ENCODE(Consortium, 2012), experiment ENCSR650QAK]. Genes (x-axis) are sorted based on their expression fold-change (calculated by limma-voom (Law et al., 2014)) between SP1-inhibited and control cells, with vertical black lines demarcating the genes that are significantly down-regulated in SP1-mutant tumors. The blue line represents the enrichment curve (Subramanian et al., 2005). (**b**) GSEA plot for SP1 over-expression in HEK293 cells (Schmitges et al., 2016). (**c**) Scatterplot of gene expression changes induced by SP1 over-expression (x-axis) and SP1 inhibition (y-axis). Genes that are significantly down-regulated in SP1-mutant tumors (mutation responsive genes) are shown with blue dots, whereas other genes are in grey. The mutation-responsive genes that are part of the leading-edge gene set in both the inhibition and over-expression GSEA analyses are highlighted. (**d**) Gene expression changes between tumor and normal tissues for SP1 (top) and its leading-edge mutation-responsive regulatory targets. Only cancer types in which SP1 is significantly up-regulated (t-test, FDR < 0.05) are shown. (**e**) Expression of leading-edge SP1 targets in tumor samples with missense mutations in SP1 ZFs (left) or missense mutations in non-ZF regions of SP1 (right). The cancer type of each ZF-mutated sample is shown on top of the heatmap. Expression values represent log fold-change relative to the average of the non-mutated samples in each cancer type, after correcting for the confounding variables. (**f-g**) Local distribution of missense polymorphisms (**f**) and somatic missense mutations (**g**) across the SP1 protein sequence. Each vertical bar represents one SNP/mutation. The curves measure the local depletion of SNPs/mutations, measured as –log10 of binomial test P-value for windows of 40 amino acids across the protein. The SP1 domain structure is shown at the bottom.

Interestingly, we found that only somatic mutations in SP1 ZFs are accompanied by significant down-regulation of the core set of SP1 targets (**Fig. 6e**). In contrast, somatic missense mutations in non-DNA-binding regions of the SP1 protein did not have a similar effect, suggesting that other mutations are not as detrimental to the regulatory function of SP1 as mutations in the ZFs. Consistent with higher deleteriousness of mutations in ZFs, we observed that SP1 ZFs are significantly depleted of missense single-nucleotide polymorphisms across the human population [based on gnomAD (Lek et al., 2016) data, binomial P < 0.001, **Fig. 6f**]. We utilized our model for domain-specific effect of SP1 mutations to test whether SP1 function is required for cancer cell function: under the hypothesis that SP1 is an oncogenic factor, this model predicts that somatic missense mutations should be depleted in SP1 ZFs relative to other SP1 regions, since such mutations would disrupt the cancer-associated regulatory function of SP1. Indeed, we observed that the ZF array of SP1, particularly ZNF1, is depleted of somatic missense mutations (binomial test, P < 0.006, **Fig. 6g**). Together, these results suggest that SP1 up-regulation in cancer promotes the expression of several cancer-associated factors, and that disruption of SP1 DNA-binding ability through mutations in its ZF array counteracts this oncogenic program, leading to negative selection of these mutations in cancer.

## DISCUSSION

Our analyses indicate that the compound recognition code (C-RC) provides a more accurate model of DNA binding by C2H2-ZFs than existing models, as a result of incorporating *in vivo* binding data during model construction. Interestingly, this improvement was achieved by inclusion of *in vivo* binding preferences of less than 840 human zinc fingers, leaving open the possibility of further improvements as additional data become available. Furthermore, our molecular dynamics analyses suggest that C-RC can quantitatively infer the contribution of different ZF residues to DNA specificity, including new potential interactions beyond the canonical model such as π-π stacking of planar amino acids with DNA bases. These interactions appear to occur as a result of alternate conformations of the amino acid side chains at the DNA-protein interface – this might explain why they were not reported earlier in the X-ray crystal structures of C2H2-ZPFs, which often represent only the most stable conformation.

By combining the C-RC with ChIP-seq and ChIP-exo data, we were able to obtain a domain-resolution map of ZFP-DNA interactions for a total of 209 ZFPs, which suggested that ∼50% of ZF domains engage with DNA *in vivo*. We observed that DNA-engaging ZFs are more conserved across species than ZF domains that do not engage with DNA. This trend in conservation, however, is not limited to the base-contacting residues and can be observed in all ZF residues, suggesting that non-base-contacting residues are also under stronger selective pressure in ZFs that engage with DNA compared to ZFs that do not engage with DNA *in vivo*. This observation is consistent with a role of non-base-contact residues in DNA-binding [e.g. through interaction with DNA backbone, as suggested by recent studies (Najafabadi et al., 2017)]. On the other hand, the relative lower sequence conservation in ZFs that do not bind DNA may reflect alternative functions that allow higher tolerance to mutations in the protein sequence.

Our functional stratification of ZFs enabled us to specifically examine the gene regulatory consequences of missense somatic mutations in DNA-binding ZFs, which revealed hundreds of ZFP-gene associations. We were able to validate these associations based on their overlap with direct regulatory targets of ZFPs, their consistence with gene expression changes that accompany loss-of-heterozygosity (LOH) at the ZFP locus, ZFP-gene co-expression across different tumours, and direct modulation of ZFPs in isogenic cell lines. However, we note that the number of genes that are directly and/or indirectly affected by the regulatory function of each ZFP is likely much higher than those we report here: First, there is a limited number of tumour samples with available transcriptomic data that harbor missense somatic mutations at DNA binding ZFs (**Fig. S6a**), which leads to a limited statistical power. Secondly, even though there is negative selection against mutations at all positions of DNA-binding ZFs, not all mutations are expected to have similar regulatory effects. Thirdly, most tumours have at most one mutated allele for each ZFP, whose loss of function might be compensated by the other, unaltered copy. For example, despite a relatively large number of mutated tumour samples with missense mutations in CTCF (38 samples), we identified only a small number of genes whose expression is affected by these mutations (35 genes, **Table S8**). Previous reports have also suggested that partial loss of CTCF by RNAi has limited regulatory consequences (Zuin et al., 2014), consistent with the limited effect of monoallelic mutations that we observed. In fact, we found that while many tumor samples have missense mutations in the DNA-binding ZFs of CTCF, these mutations are mutually exclusive with loss-of-heterozygosity (LOH) at the CTCF locus (Fisher’s exact test P < 3×10^−4^, **Fig. S6b**), suggesting a selective pressure in cancer against complete loss of CTCF function.

Interestingly, our pan-cancer analysis also revealed a substantial number of genes whose expression is affected by somatic mutations in the DNA-binding ZFs of KRAB proteins (**Fig. 5c**), which are mostly studied in the context of retroelement repression (Thomas and Schneider, 2011) and whose functions in regulating host gene expression are less understood. This observation suggests that mutations in these proteins have widespread consequences on the transcriptome, which is consistent with more recent views of regulation of host genes by KRAB proteins in addition to their retroelement-inhibitory role (Imbeault et al., 2017; Najafabadi et al., 2015b). The regulatory role of KRAB proteins is also supported by the higher conservation of DNA-binding ZFs in these proteins and their depletion from common genetic variants at the population level, suggesting that mutations in the DNA-binding domains of KRAB proteins are deleterious. This is a surprising finding given that there are few known disease-linked mutations in KRAB proteins, and highlights the need for better characterization of the functions of these proteins in genome regulation.

In this regard, a question that needs to be addressed is whether the KRAB proteins that we analyzed are representative of all KRAB-containing ZFPs: We limited our analyses to the ZFP binding sites that do not overlap with endogenous repeat elements (EREs) due to their confounding effect on motif finding (Najafabadi et al., 2015a); however, a large number of the KRAB protein binding sites are in EREs (Imbeault et al., 2017; Najafabadi et al., 2015b; Schmitges et al., 2016), which may result in the identification of high-quality motifs for fewer KRAB ZFPs. Indeed, we were able to obtain high-quality motifs for only 25% of the KRAB proteins that almost exclusively bind to ERE regions (i.e. >90% of top 500 peaks in EREs). In contrast, for the rest of KRAB proteins, we had >97% success rate in finding high-quality motifs that match the recognition code predictions, suggesting that our results may be biased against exclusive ERE binders. We note, however, that this group likely forms a relative minority of KRAB proteins (12 out of 55 KRAB proteins with ChIP-seq data match our definition of exclusive ERE binder).

Our study of the *in vivo* DNA-binding ability of ZFs mirrors previous studies that have shown that, at least *in vitro* and when fused to ZF1-2 of the Egr1 protein, not all ZF domains are able to bind to DNA (Najafabadi et al., 2015b). These studies suggest that there are intrinsic differences between DNA-binding and non-binding ZFs. Do these intrinsic differences explain the observation that only a fraction of human ZF domains engage with DNA *in vivo*? We have found that ZFs that bind to DNA *in vivo* have, on average, higher predicted sequence specificity than ZFs that do not engage with DNA (**Fig. S5d**), as predicted by our recognition code. However, sequence specificity alone is a modest predictor of *in vivo* DNA binding. Furthermore, ZFs that do not engage with DNA *in vivo* have, on average, significantly higher sequence specificity than ZFs in pseudogenes (which are presumably non-functional), suggesting that at least some of the non-engaging ZFs are intrinsically capable of binding to DNA (**Fig. S5d**). This notion is also supported by direct comparison of our *in vivo* binding map with *in vitro* bacterial one-hybrid data (Najafabadi et al., 2015b), which shows that while there is a statistically significant overlap between ZFs that bind to DNA *in vitro* and *in vivo* (odds ratio=1.7, P < 0.003, Fisher’s exact test), at least 15% of ZFs that do not engage with DNA *in vivo* are still capable of binding to DNA *in vitro* (**Fig. S5e**). This raises the possibility that DNA-binding capability is, at least to some extent, determined by the context, which may include the ZF position in the array as well as the identity of adjacent ZFs. More strikingly, ∼55% of ZFs that do not bind DNA *in vitro* interact with DNA *in vivo*, often with high specificity, enforcing the notion that *in vitro* binding assays at best only partially reflect the *in vivo* function of ZFs.

Finally, we note the possibility that some ZFs may engage with DNA in alternative binding modes that we have not yet characterized. A few ZFPs are known to employ alternative sets of ZFs to bind to DNA (Baxley et al., 2017; Han et al., 2016; Nakahashi et al., 2013). This observation suggests that some ZFs might be engaged only in a subset of *in vivo* binding sites – such ZFs would not be part of the core motifs that we have presented in this paper, which can potentially explain their higher-than-random sequence specificity. Therefore, the factors that determine the ability of ZFs to engage with DNA may depend not only on the ZFP itself, but also on its binding sites.

## METHODS

### Obtaining the data set of C2H2-ZF preferences for training the recognition code

We obtained the *in vivo* binding sites of 313 proteins from two previous studies (Imbeault et al., 2017; Schmitges et al., 2016), representing 131 proteins with ChIP-seq and 221 proteins with ChIP-exo data in HEK293 cells. The binding sites of each protein were directly downloaded from GEO datasets associated with these publications (GEO accessions GSE76494 and GSE78099). For each datasets, we ran RCADE (Najafabadi et al., 2015a) on the top 500 peaks that did not overlap any endogenous repeat elements (EREs), as described previously (Dogan and Najafabadi, 2018) [ERE coordinates were obtained from the RepeatMaser track of the UCSC Genome Browser(Karolchik et al., 2004)]. For successful runs, we included the top-ranking motif of each protein in our training dataset. For the proteins that had both ChIP-seq and ChIP-exo data and resulted in a significant motif in both cases, we used the ChIP-exo motif, given the higher resolution of ChIP-exo compared to ChIP-seq. We split each motif into its constituent triplets (total of 836 triplets), and also extracted the ZF associated with each triplet, as well as the two ZFs on the two ends of each direct ZF. For the triplet at the 5’ end of each motif, no ZF from the C-terminus of the associated ZF was used. Similarly, for the triplet at the 3’ end of each motif, no ZF from the N-terminus of the associated ZF was used (see **Fig. S2** for a schematic representation). We set aside 50 randomly selected ZFs from this training dataset, so that they could be used at the end for testing the obtained model. We augmented the training dataset with *in vitro* B1H-based motifs from 8138 ZFs (Najafabadi et al., 2015b). Since in these B1H experiments only the binding preference of a single ZF was queried at a time, the associated ZF of each motif does not have any N-terminal or C-terminal adjacent ZFs.

### Biochemical encoding of amino acids

For each amino acid, we extracted six different biochemical properties (Yousef and Charkari, 2015) (**Table S3a**). Given the strong correlations among these properties, we used principal component analysis (PCA) and used the transformed coordinates of the amino acids on the first two components (**Table S3b**) for downstream analyses.

### Feature selection and training of the recognition code

We considered four possible input configurations for predicting the preference of the ZF for a DNA triplet: (a) only the ZF that is directly in contact with the triplet, (b) the direct ZF plus its N-terminal neighbor, (c) the direct ZF plus its C-terminal neighbor, and (d) the direct ZF plus its two neighbors.

For each of these configurations, we considered different sets of input features (**Figure S2b**): (i) only the four canonical residues of the ZFs (i.e. residues –1, +2, +3, and +6), (ii) the seven residues that showed the highest correlations with the DNA preference according to Chi-square test of *in vivo* data (i.e. residues –4, –2, –1, +1, +2, +3, and +6), and (iii) all 12 residues between the second Cys and the first His in the ZF (i.e. the X_12_ in the Pfam pattern X_2_CX_2,4_CX_12_HX_3,4,5_H). For each configuration a-d, we tried each feature set i-iii for predicting each of the four bases at each position of the triplet, and chose the feature set that maximized the Pearson correlation of predicted vs. target values during 5-fold cross-validation.

Finally, for each of the three possible ZF contexts, we compared the performances of the compatible configurations and selected the best-performing configuration by 5-fold cross-validation. The ZF contexts were: (1) ZFs that are at the N-terminus of the DNA-binding ZF array (compatible with configurations a and c), (2) ZFs that are at the C-terminus of the DNA-binding array (compatible with configurations a and b), and (3) ZFs that in the middle of the DNA-binding array (compatible with configurations a, b, c, and d). The configuration that was selected for each context and the corresponding feature set are shown in **Fig. S2c**.

### Evaluating the performance of the final mode

After configuration- and feature-selection based on 5-fold cross-validation results, the final recognition code was tested on 50 randomly selected held-out ZFs that were not included at any stage of training. We used the Pearson correlation of predicted vs. observed base probabilities as the measure of performance. Also, we used MoSBAT-e (Lambert et al., 2016) to compare the predictions of the code with motifs obtained from protein binding microarray analysis of a set of non-animal ZFPs (Najafabadi et al., 2017) or chimeric ZFPs constructed from fusion of different ZFs (Lam et al., 2011).

### Molecular dynamics simulations of Egr1 and CTCF

CTCF-DNA complex was obtained by combining two crystal structures (PDB accessions 5T0U, 5UND) (Hashimoto et al., 2017), and EGR1-DNA complex was obtained from PDB accession 4×9J (Zandarashvili et al., 2015). The simulations were carried out using AMBER16 package (Case et al., 2005) with the *ff14SB* force field. The system was immersed in a rectangular box of TIP3P water model (Jorgensen et al., 1983) with neutralizing concentration of ions. Additionally, Na+/Cl–ions matching the ionic strength of 0.1 M were included. Each system was energy-minimized using 2500 steps of steepest descent followed by 2500 steps of conjugated gradient using a harmonic restriction on the solute with a value of 40 kcal/mol·Å. The heating process was carried from 0 to 300 K using Langevin dynamics and subsequently followed by equilibration for 500 ps (NPT). Finally, we carried out unrestrained MD for 1 μs under NPT conditions. The particle mesh Ewald (PME) method (Essmann et al., 1995; Galindo-Murillo et al., 2013) was used to handle long-range electrostatic interactions and a cutoff of 12.0 Å was used for the simulations. SHAKE algorithm(Ryckaert et al., 1977) was used to contain hydrogen bonds and to allow a longer integration step. All simulations were carried out with a time step of 2 fs using the *pmemd* module in AMBER16 (Salomon-Ferrer et al., 2013). The *cpptraj* module was used for trajectory analyses (Roe and Cheatham, 2013). The criterion for hydrogen bonding was set at ≤3.0 Å distance between electron donor atom and hydrogen of electron acceptor atom with 120-degree angle cutoff. Heavy atoms involved in stacking interaction were identified as described in MolBridge (Kumar et al., 2014). A perpendicular distance cutoff of 3.2 Å between the planar atoms was used. The simulated structure and contacts were visualized using PyMOL (Schrodinger, 2015).

### Identification of the *vivo* binding preferences and DNA-engaging domains of human ZFPs

We created a modified version of RCADE (Najafabadi et al., 2015a), called RCADE2, which uses our new compound recognition code (C-RC) to predict the binding preferences of different ZF arrays, evaluate their enrichment in *in vivo* binding sequences (compared to dinucleotide-shuffled sequences), and then optimize the motifs to maximize AUROC. RCADE2 is available at https://github.com/csglab/RCADE2, and includes features such as HTML output and integrated positional analysis of the identified motifs. Using RCADE2, we analyzed ChIP-seq and ChIP-exo data from two previous publications (Imbeault et al., 2017; Schmitges et al., 2016) as described above, including removal of peaks that overlapped EREs to ensure that our analyses are not confounded by sequence homology among repeat elements, and then using the summit position of the top 500 peaks with the largest scores (i.e. smallest p-values) for motif finding. For successful runs, we extracted the top-scoring optimized motif that had significant similarity with the seed motif (i.e. the motif predicted by C-RC) at P < 0.001, and marked the ZFs associated with that motif as the DNA-engaging ZFs. For each DNA-engaging ZF, we also calculated the information content of the 3-nucleotide motif that it encodes as *IC* = ∑_*i,j*_ *p*_*i,j*_*log*_2_(4 × *p*_*i,j*_), where *p*_*i,j*_ denotes the probability of observing base *j* at position *i* of the triplet motif (1≤*i*≤3).

### Analysis of cross-species conservation and population-level genetic variation at ZFs

We obtained missense single-nucleotide variants (SNVs) and their frequencies from the Genome Aggregation Database, gnomAD (Lek et al., 2016), for genomic build GRCh37.24. Only single nucleotide variants that passed the high-quality filter in gnomAD (PASS filter) were included for further analysis. Multiple variants for the same genomic location were split using BCFTools. ZF domains were annotated based on the presence of Pfam pattern X_2_CX_2,4_CX_12_HX_3,4,5_H in the protein sequence, and the SNVs from the canonical transcript were mapped to their corresponding amino acid residue in the ZF. To obtain per-residue genetic variation, allele frequencies of the three codon positions were summed. We used a similar approach to analyze per-base conservation at the ZF coding sequences, using phyloP (Pollard et al., 2010) conservation scores that were obtained from the UCSC Genome Browser (phyloP46way track, hg19).

### Identification of ZFP-gene associations based on ChIP-seq data

To identify genes that have a binding site for each ZFP near their TSS regions, we first identified the high-confidence set of peaks for each ZFP by simultaneous optimization of the peak-calling score cutoff and the motif hit score cutoff, similar to a previous approach (Schmitges et al., 2016). Briefly, for each ZFP, we first identified the affinity-based motif-match score of each peak using AffiMx (Lambert et al., 2016). We then identified a motif score cutoff to dichotomize the peaks, in a way that maximizes the Mann-Whitney U z-statistic of the difference of peak scores between the two peak sets. Similarly, we identified a peak score threshold that maximizes the Mann-Whitney U z-statistic of the difference of motif scores between peaks that pass that threshold and peaks that do not. Any peak that passed the optimized peak-score threshold was deemed a significant binding site, and any significant binding site that also passed the optimized motif-score threshold was deemed to have a match to the motif. Then, we associated a gene to a significant binding site if that binding site was within 10kb of the TSS of the gene (**Data File S4**).

### Data for mutation/expression analysis across cancers

Somatic mutations, copy number alterations (CNA), patient metadata (including age and sex), and gene expression data of TCGA Pan-Cancer Atlas (Hoadley et al., 2018) were downloaded from cBioPortal (Gao et al., 2013). Gene expression values were normalized by division to the 75^th^ percentile of the non-zero values of each sample. We retained only tumour samples that had all of the four data types mentioned above. We further filtered the samples to keep only those that had at least one missense mutation in at least one of the ZFPs with ChIP-seq data that we studied here. We also removed datasets (tissues) for which there were fewer than 30 samples with at least one mutated ZFP. These filters resulted in a total of 2538 tumour samples from 18 cancer types (**Table S6**). Missense mutations were then classified into those that overlap a DNA-binding ZF (i.e. ZFs that recognize, *in vivo*, a 3-nucleotide motif with IC ≥ 1; see previous sections), those that overlap a non-DNA-binding ZF, and those that do not overlap a ZF. A total of 96 ZFPs had at least one mutation overlapping a DNA-binding ZF, which were further analyzed for association of their mutations with genome-wide expression.

### Identification of genes whose expression is associated with ZFP mutations

Gene expression data were first filtered to include only genes that had an expression value >0.01 in at least 25% of all samples. For each ZFP *z*, we then selected the samples that did not have a CNA at the ZFP locus (i.e. CNA=0 based on the cBioPortal data), and further normalized and log-transformed the expression values using voom (Law et al., 2014). Then, for each gene *i*, we fit a linear model to the normalized and log-transformed expression data across samples *j* in the form of *E*_*i*_∼*T*+*T*×*S*+*T*×*A*+*N*_*i*_+*M*_*z*_, where *E*_*i*_ is the vector of expression of gene *i* across samples, *T* is the vector of tissue of origin of the samples, *S* denotes the sex of the patients from which the samples were obtained, *A* represents the age of the patients, *N*_*i*_ denotes the CNA status of gene *i* across samples (with integer values ranging from –2 to +2), and *M*_*z*_ is the vector of mutation status of the ZFP *z* across samples (non-mutated, mutation in a non-ZF region, mutation in a DNA-binding ZF, and mutation in a non-DNA-binding ZF). The coefficients of the variable *M*_*z*_ for DNA-binding ZF mutations, along with the associated P-values, were then used to identify genes whose expression is associated with DNA-binding-disrupting mutations. Overall, this approach is similar to a previous work on identification of *cis*-eQTLs based on somatic mutations (Zhang et al., 2018), with the difference that we looked at *trans*-acting coding mutations, we aggregated the mutations by the function of the affected protein domain, and we did not include any hidden factors in the regression model – inclusion of hidden factors are necessary to correct for the global structure and correlations in gene expression data in order to identify *cis*-acting effects; however, gene-gene correlations are likely driven by *trans*-acting factors, and therefore we decided not to remove them in our *trans*-eQTL analysis.

### Overlap of ChIP-seq-based regulatory links and expression-mutation associations

To examine whether expression-mutation associations represent regulatory relationships between ZFPs and genes, we calculated enrichment of ChIP-seq-based ZFP-gene regulatory links, as determined by presence of a ChIP-seq binding site within 10kb of TSS (see above). For **Fig. 4c-f**, we examined the enrichment of these regulatory links in ZFP-gene pairs that fall within a specific bin (binned by the effect size of DNA-binding ZF somatic mutations in **Fig. 4c** and **e**, and by the joint distribution of the effect size of over-expression and somatic mutations in **Fig. 4d**). For each bin, we then performed a logistic regression in order to model the log-odds of presence of a regulatory link as a function of the bin in the form of *L∼B*, where *L* is the log-odds that a ZFP-gene pair corresponds to a ChIP-seq-based regulatory link, and *B* is a binary vector indicating whether that ZFP-gene pair belongs to the bin that is being examined. We note that some genes have overall higher probability of being associated with at least one ZFP in our ChIP-seq compendium, which may represent a bias towards genes that are expressed in HEK293 cells (Schmitges et al., 2016). To correct for this bias, in the logistic regression, we considered the presence of a regulatory link between the ZFP and the gene as a “success”, and the number of ChIP-seq-based regulatory links that connect that gene to “other” ZFPs as the number of “failures”. Therefore, for example, if gene *x* has at least a peak for ZFP *z*, and also at least one peak for *n* other ZFPs, then the regression assumes *n+1* Bernoulli observations for the *x*-*z* pairs, in which 1 was a success and *n* were failure. On the other hand, if gene *x* didn’t have a peak for ZFP *z*, then there would be 0 success and *n* failures. We used a similar logistic regression for **Fig. 4f**, with the difference that the predictor variable, instead of being a binary vector, was the ZFP mutation effect size.

### Identification of genes whose expression is associated with loss-of-heterozygosity (LOH) of ZFPs

We performed another association analysis, similar to the regression approach mentioned above, to identify genes whose expression is affected by LOH of each ZFP. Specifically, for each ZFP *z*, we excluded all samples that had any mutations in the ZFP coding sequence as well as samples that had biallelic deletion or copy number gains at the ZFP locus, normalized and log-transformed the resulting expression matrix using voom (Law et al., 2014), and then fit a model of the form *E*_*i*_∼*T*+*T*×*S*+*T*×*A*+*N*_*i*_+*N*_*z*_, where *N*_*z*_ is the binary vector of CNAs for ZFP *z* across the samples (0 for no CNA, 1 for LOH).

### Identification of significant associations between gene expression and somatic mutations

We observed that LOH-associated gene expression changes are highly consistent with gene expression changes that are associated with mutations in DNA-binding ZFs. Therefore, we used LOH association to form a prior for P-value adjustment and detection of significant mutation associations. Specifically, we used independent hypothesis weighting, IHW (Ignatiadis et al., 2016), to give a weight to each ZFP-gene pair based on the t-score of the coefficient of *N*_*z*_ in the LOH-expression association analysis, and then use these weights to maximize statistical power for detection of significant mutation-expression associations at FDR < 0.2.

### Concordance between gene signatures of each ZFP and ZFP-gene co-expression

For each ZFP *z* and each cancer type, we modeled the expression of each gene *i* as a linear function of the expression of the ZFP across samples, taking into account the confounding effects of age and sex and the CNAs at the query gene locus: *E*_*i*_∼*S*+*A*+*N*_*i*_+*E*_*z*_, where *E*_*z*_ is the vector of expression of ZFP *z* across tumours. Then, for each ZFP, we calculated the Pearson correlation between the coefficients of *E*_*z*_ across the genes that are part of the ZFP regulon (based on association with ZFP mutation, see above) and the ZFP mutation effect size. Note that since the effect of somatic mutations in DNA-binding ZFs is expected to be negative for co-expressed ZFP-gene pairs and positive for anti-correlated ZFP-gene pairs, the Pearson correlation sign is inverted.

## Supporting information

Supplementary Data File S1 (ChIP-seq motifs)

Supplementary Data File S2 (ChIP-exo motifs)

## ACKNOWLEDGEMENTS

This work was supported by funds from the Natural Sciences and Engineering Research Council of Canada (RGPIN-2018-05962), Canadian Institutes of Health Research (PJT-155966), and resource allocations from Compute Canada to H.S.N. H.S.N holds a Canada Research Chair funded by the Canadian Institutes of Health Research. B.D. is supported by a postdoctoral fellowship from the Scientific and Technological Research Council of Turkey (TUBITAK, grant no. 1059B191600423). A.H.C. is supported by a Globalink Graduate Fellowship from Mitacs.

## AUTHOR CONTRIBUTIONS

B.D. and H.S.N. developed the computational methods and analyzed the data. S.K. performed the structural analyses and contributed to analysis of genetic variations. A.H.C. contributed to data analysis and algorithm implementation. N.K. contributed to collection and processing of cancer data. H.S.N. conceived and directed the study. H.S.N wrote the manuscript with contribution from B.D. and S.K.

