## Supplementary Figures for "A domain-resolution map of *in vivo* DNA binding reveals the regulatory consequences of somatic mutations in zinc finger transcription factors"

[1] Department of Human Genetics, McGill University, Montreal, QC, Canada
[2] McGill Genome Centre, Montreal, QC, Canada
[3] Rosell Institute for Microbiome and Probiotics, Montreal, QC, Canada
[4] Current address: Department of Biomedical Engineering, Inonu University, Malatya, Turkey

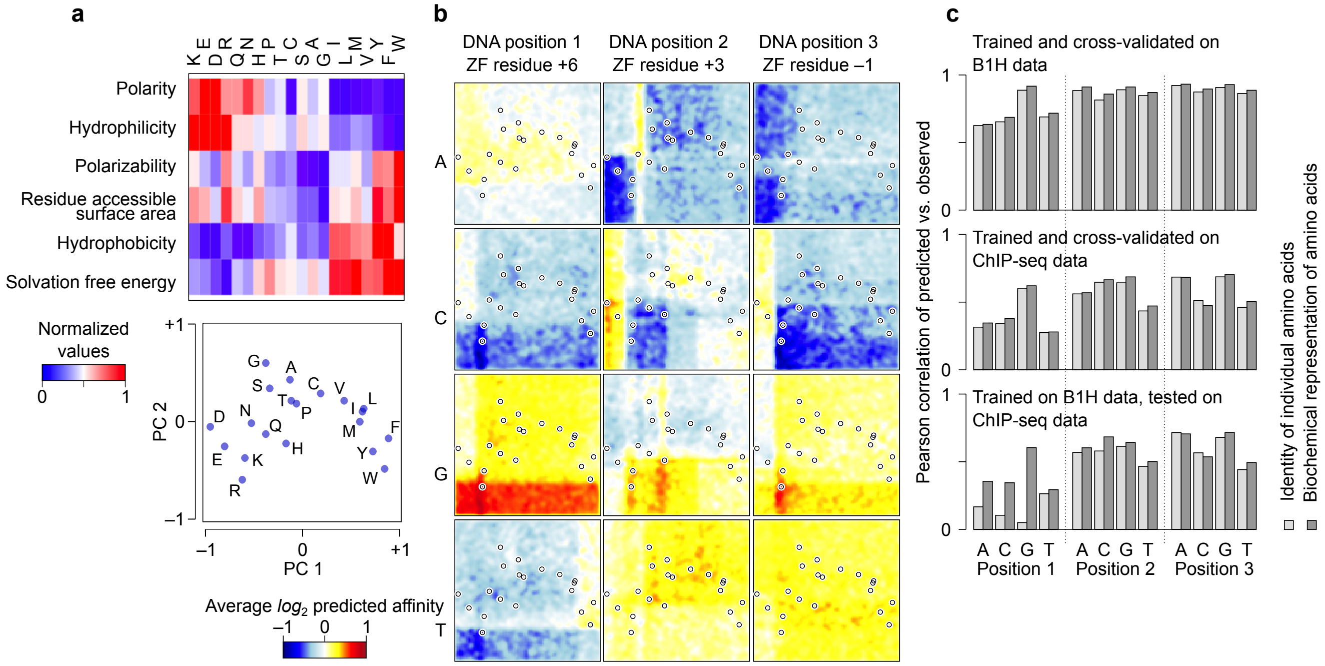


**Figure S1. Encoding the amino acids by their biochemical properties.** (**a**) The heatmap (top) represents the biochemical properties we considered. The PCA-transformed values are shown in the scatterplot at the bottom. Underlying data are provided in **Table S3**. (**b**) Encoding the amino acids based on their PCA representation allows a random forest regression model to learn simpler rules that are shared among amino acids with similar properties. The figure shows a visual representation of example rules learned by random forest. Specifically, we generated 40,000 "pseudo-amino acids" by dividing the PCA plot in panel (a) into a 200×200 grid, and then generated 40,000 random ZFs by sampling (without replacement) these pseudo-amino-acids for each of the 12 ZF positions. The color gradient in the graphs shows the predicted affinity of these random ZFs for recognition of each base at each position of the DNA triplet. The ZFs are projected on the scatterplot based on the PCA coordinates of the pseudo-amino-acid at position +6, +3, or –1, as indicated above the graph. (**c**) The performance of the recognition code for predicting the probability of each base at each triplet position, when the amino acids are encoded as categorical variables (20 individual identities) or using the PCA-transformed biochemical properties. We used 5-fold cross-validation on B1H motifs[3] (top) or ChIP-seq motifs[1] (middle), or trained the recognition code on B1H motifs and tested on ChIP-seq motifs (bottom). For the latter, we ensured that any ZF present in the ChIP-seq data was removed from the B1H training set to prevent the over-estimation of performance.

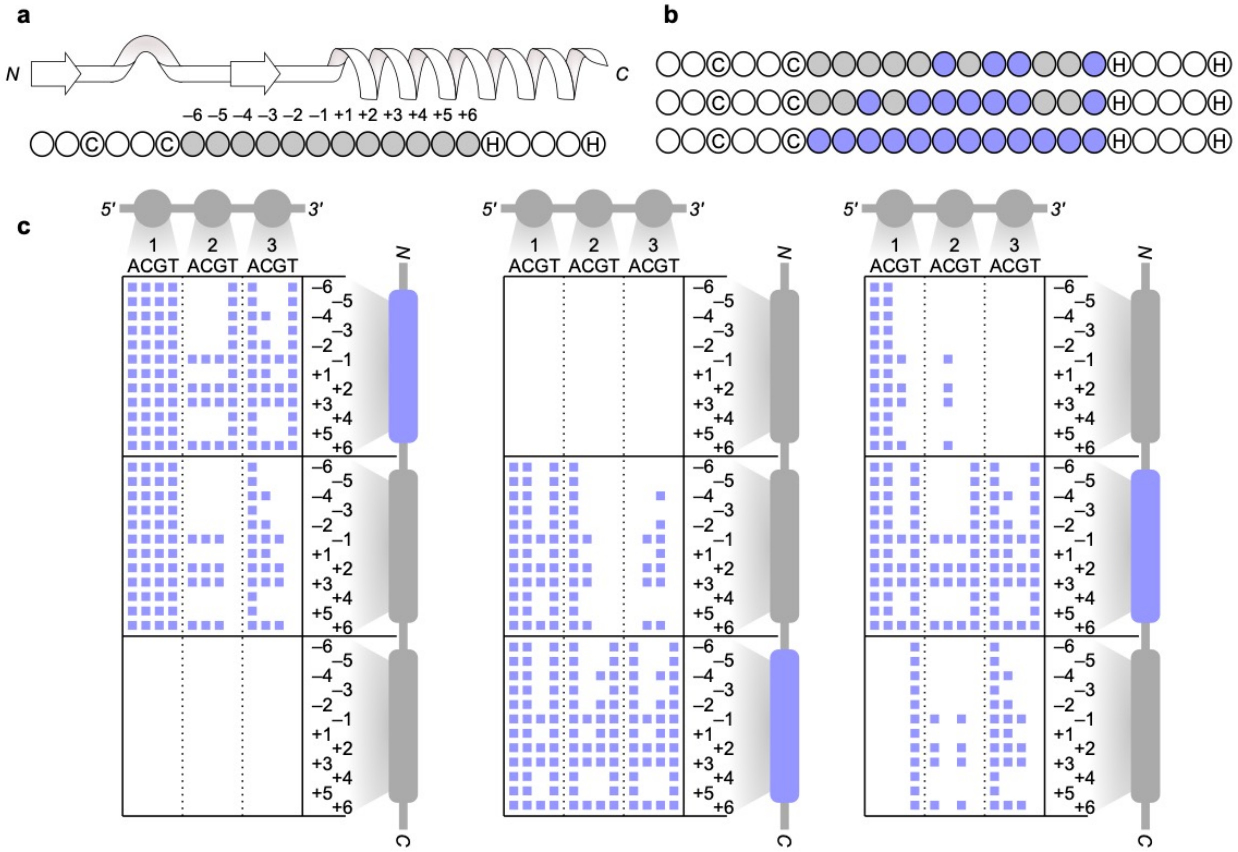


**Figure S2. Features used by the compound recognition code (C-RC).** (**a**) The structure and residue numbering in C2H2-ZFs. Arrows, loop, and helix represent the beta-sheets, turn, and alpha helix structures, respectively. (**b**) The different sets of residues that were considered as input features for training C-RC: the four canonical residues of the ZFs (top), the seven residues that showed the highest correlations with the DNA preference according to Chi-square test of *in vivo* data (middle), and all 12 residues between the second Cys and the first His in the ZF (bottom). In each case, the residues that are used as predictor variables are highlighted in blue. (**c**) The optimal feature sets that were selected for each of the 36 random forests (4 bases x 3 triplet positions x 3 ZF contexts). Each panel shows one of the three ZF contexts: predicting the DNA triplet that is adjacent to an N-terminal ZF (left), C-terminal ZF (middle), or non-terminal ZF (right). In each panel, each column corresponds to the random forest model for predicting the preference for one base at one of the triplet positions, and blue squares represent the features that are included in each random forest model.

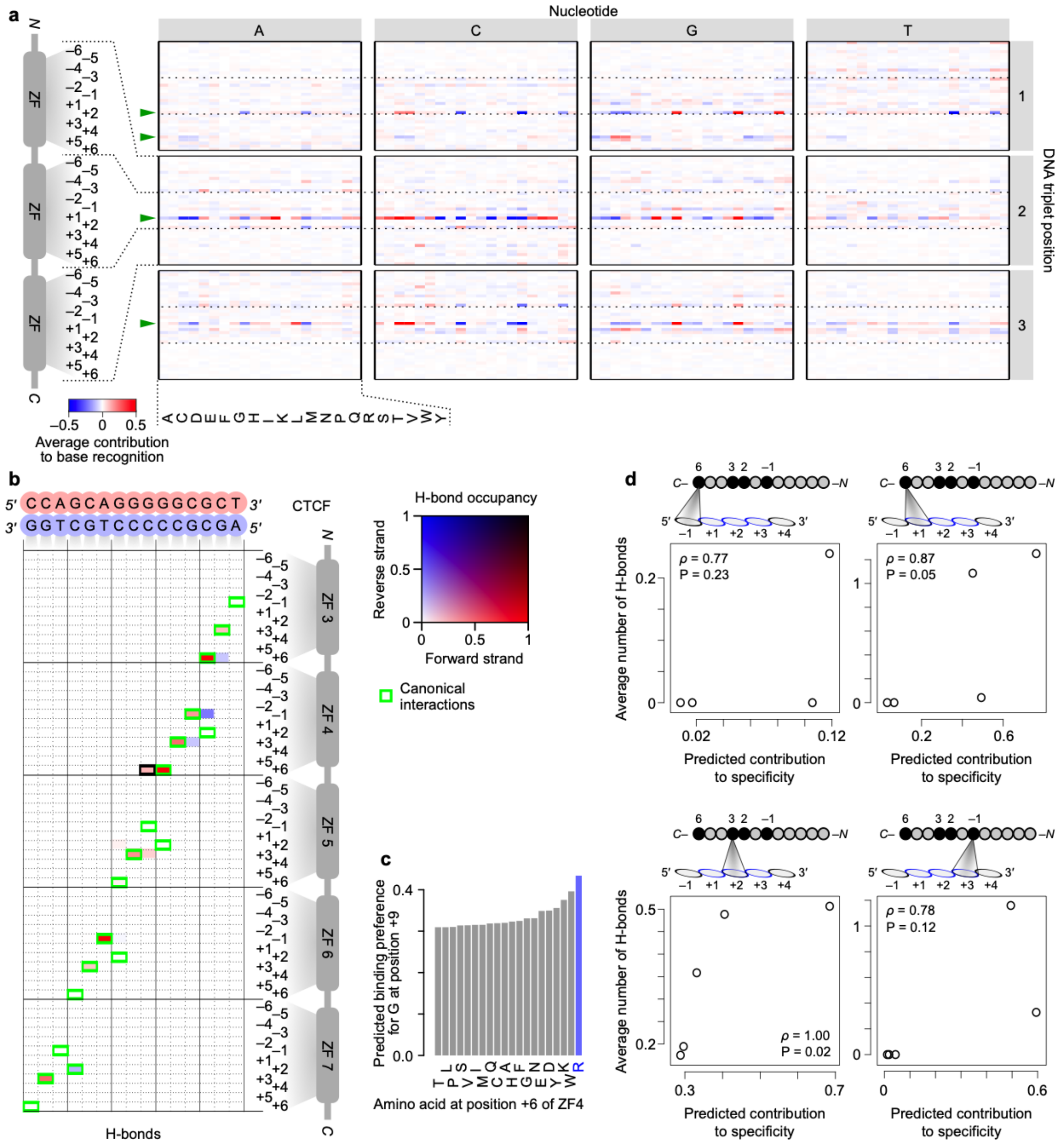


**Figure S3. C-RC quantitatively predicts amino acid-base interactions.** (**a**) To understand how C-RC works, we examined the output of the code for 100,000 randomly generated ZFs, each with one randomly generated adjacent ZF on each side. We then used linear regression to examine the association of the output of C-RC with the amino acid identities at each of the central or adjacent ZF positions. The heatmap shows the regression coefficients, representing the average contribution of each amino acid at different ZF positions for recognition of each base at different DNA triplet positions. Each panel represents recognition of one base, indicated above the figure, in a specific DNA triplet position, indicated on the right; each column represents one amino acid, each row represents one ZF position, and the color gradient denotes the contribution toward specificity (red: increased preference for the specified base; blue: decreased preference). The dashed lines separate three consecutive ZFs, where the DNA triplet would be directly adjacent to the middle ZF. The specificity residues that, according to the canonical model, contribute to the recognition of each DNA position are shown with green arrows. (**b**) H-bonds identified from MD simulations of CTCF in complex with its target DNA. The interactions in the canonical C2H2-ZF model are shown with a green border. The non-canonical interaction highlighted in panel b is shown here with black border. Underlying data are provided in **Table S4**. (**c**) The predicted preference of variants at position +6 of ZF4 for binding to G at DNA position 9 (i.e. position –1 relative to the ZF4-associated triplet). The wild-type amino acid is highlighted in blue. (**d**) Scatterplots for the recognition code-predicted associations vs. MD simulation-based H-bonds. In each plot, each dot represents one ZF. The graphs represent the amino acid-base pairs for which at least one H-bond and at least one non-zero association based on the recognition code was found. The spearman correlations are shown, along with their associated P-values (two-tailed).

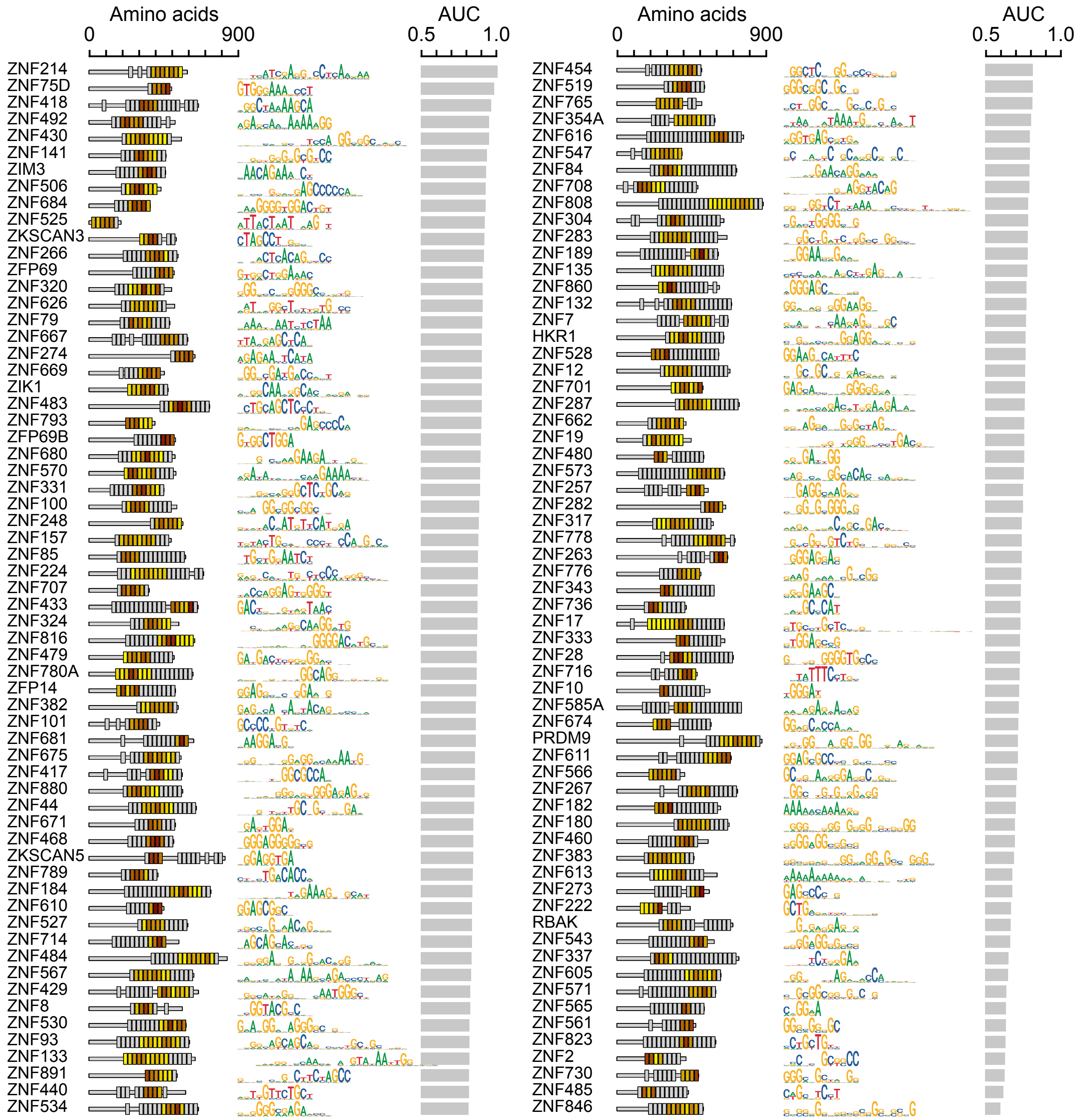


**Figure S4. Motifs identified by recognition code-assisted analysis of ChIP-exo[2] data for C2H2-ZFPs.** Annotations are similar to **Fig. 3a**.

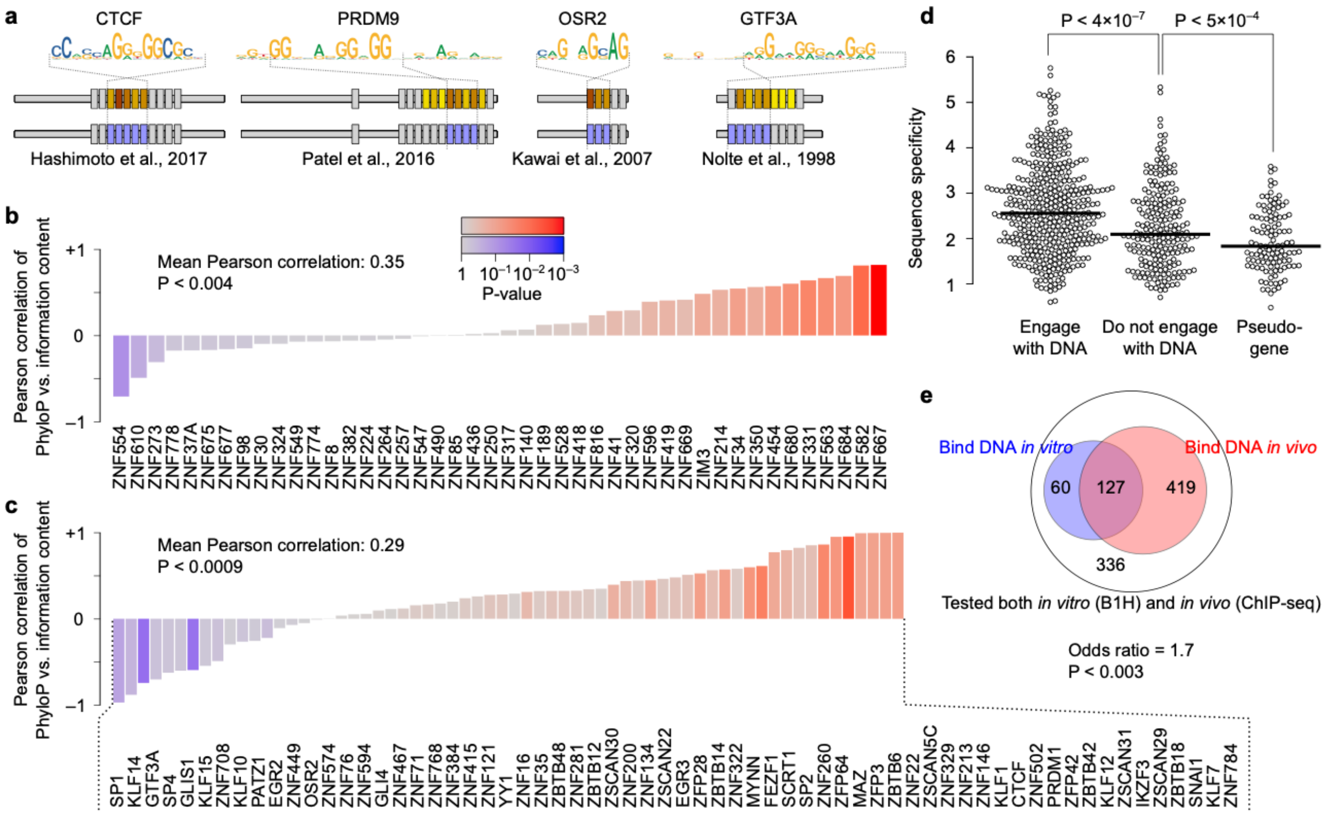


**Figure S5. Association of *in vivo* DNA binding with conservation and sequence specificity.** (**a**) Consistence between C-RC-predicted DNA-binding ZFs and previously reported DNA-binding ZFs for four C2H2-ZFPs. In each case, the optimized *in vivo* motif is shown on top, and the corresponding domains based on C-RC predictions are shown in the middle (with the color gradient showing the information content of the recognized triplet as in **Fig. 3a**) The DNA-interacting ZFs based on previous studies is shown at the bottom (DNA-interacting ZFs are in blue). Previously reported DNA-interacting ZFs: CTCF: ref[4], PRDM9: ref[5], OSR2: ref[6], GTF3A: ref[7]. (**b**) Pearson correlation of the phyloP score with the information content (IC) of the triplet recognized by each ZF. Only KRAB proteins are included in the bar plot. Annotations are similar to **Fig. 3c**. (**c**) Pearson correlation of phyloP vs. IC for non-KRAB proteins (similar to panel b). (**d**) Association between *in vivo* DNA binding and sequence specificity. We defined sequence specificity of each ZF as the logarithm of the ratio of the probabilities of binding to the most-preferred triplet vs. binding to the least-preferred triplet, as predicted by C-RC. (**e**) Venn-diagram of the overlap between ZFs that were previously reported to bind to DNA *in vitro*[3] and those that we found, based on analysis of ChIP-seq data[1], to engage with DNA *in vivo*.

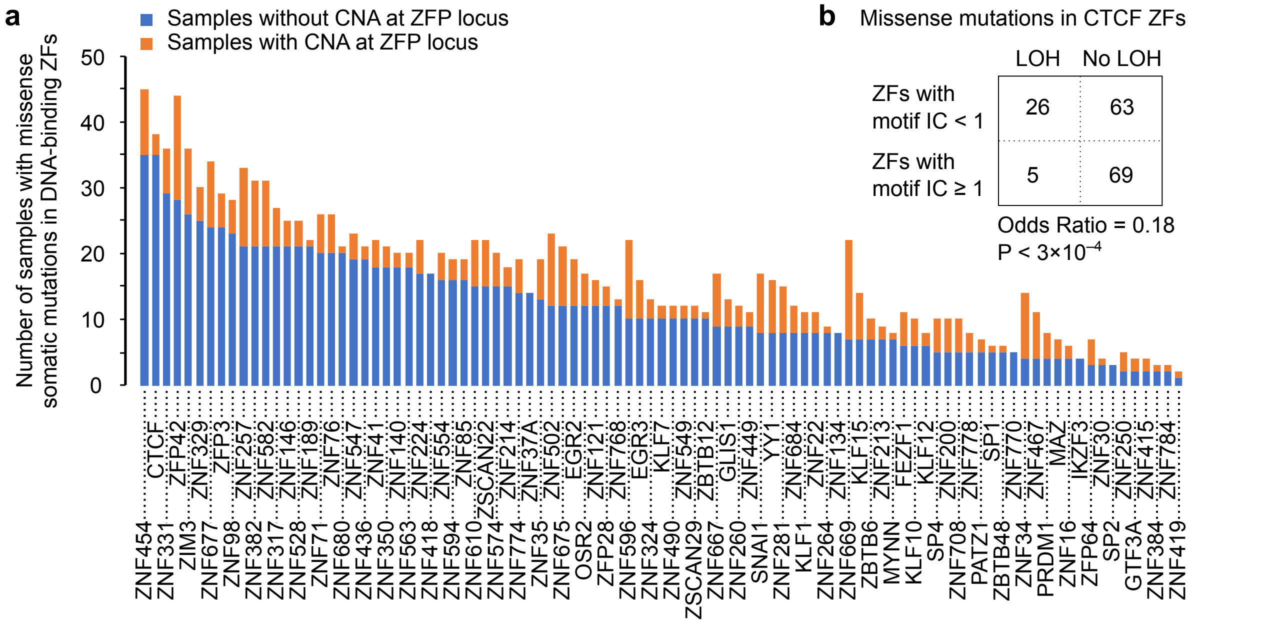


**Figure S6.** (**a**) Summary of TCGA samples with ZF somatic mutations. Note that only samples with no copy number alterations (CNAs) at each ZFP locus were used for identification of gene-ZFP associations. (**b**) Contingency table showing the number of samples with somatic mutation in DNA-binding ZFs of CTCF (ZFs with information content or IC ≥ 1 bit) and non-DNA-binding ZFs of CTCF (IC < 1) stratified by loss-of-heterozygosity (LOH) at CTCF locus. Note that this table corresponds to the curated non-redundant set of tumors from cBioPortal[8], which do not necessary have associated RNA-seq data. Therefore, the sample numbers in this table are larger than those in panel (a) which was limited to the TCGA samples with RNA-seq data.
